# All-trans retinoic acid-mediated ADAR1 degradation synergizes with PD-1 blockade to suppress pancreatic cancer

**DOI:** 10.1101/2024.10.20.619300

**Authors:** Ching-Fei Li, Li-Yuan Bai, Yongkun Wei, Heng-Huan Lee, Riyao Yang, Jun Yao, Huamin Wang, Ying-Nai Wang, Wei-Chao Chang, Yi-Chun Shen, Shao-Chun Wang, Cheng-Wei Chou, Jie Fu, Jianhua Ling, Yu-Yi Chu, Chang-Fang Chiu, Michael Wang, Dihua Yu, Paul J Chiao, Han Liang, Anirban Maitra, Haoqiang Ying, Mien-Chie Hung

**Affiliations:** Department of Molecular and Cellular Oncology, The University of Texas MD Anderson Cancer Center; Houston, TX 77030, USA; Division of Hematology and Oncology, Department of Internal Medicine, China Medical University Hospital, and China Medical University; Taichung 40402, Taiwan; Department of Lymphoma and Myeloma, The University of Texas MD Anderson Cancer Center; Houston, TX 77030, USA; Department of Pathology. The University of Texas MD Anderson Cancer Center; Houston, TX 77030, USA; Center for Molecular Medicine, China Medical University Hospital, China Medical University; Taichung 40447, Taiwan; Graduate Institute of Biomedical Sciences, College of Medicine, China Medical University; Taichung 40402, Taiwan; Research Center for Cancer Biology, China Medical University; Taichung 40402, Taiwan; Department of Biotechnology, Asia University; Taichung 41354, Taiwan; Cancer Center, China Medical University Hospital, China Medical University; Taichung 404332, Taiwan; The University of Texas MD Anderson UTHealth Graduate School of Biomedical Sciences; Houston, TX 77030, USA; Department of Bioinformatics and Computational Biology, The University of Texas MD Anderson Cancer Center, Houston, TX 77030, USA; Graduate Program in Quantitative and Computational Biosciences, Baylor College of Medicine, Houston, TX 77030, USA; Department of Systems Biology, The University of Texas MD Anderson Cancer Center, Houston, TX 77030, USA; Department of Translational Molecular Pathology, The University of Texas MD Anderson Cancer Center, Houston, TX 77030, USA; Sheikh Ahmed Center for Pancreatic Cancer Research, The University of Texas MD Anderson Cancer Center, Houston, TX 77030, USA

## Abstract

As a double-stranded RNA–editing enzyme and an interferon-stimulated gene, double-stranded RNA-specific adenosine deaminase (ADAR1) suppresses interferon signaling and contributes to immunotherapy resistance. Suppression of ADAR1 overcomes immunotherapy resistance in preclinical models, but has not yet been translated to clinical settings. By conducting a screening of a subset of the FDA-approved drugs, we found that all-trans retinoic acid (ATRA, also known as tretinoin) caused ADAR1 protein degradation through ubiquitin-proteasome pathways and concomitantly increased PD-L1 expression in pancreatic and breast cancers. In addition, the combination of ATRA and PD-1 blockade reprogrammed the tumor microenvironment and unleashed antitumor immunity and thereby impeded tumor growth in pancreatic cancer mouse models. In a pilot clinical trial, a higher dose of ATRA plus the anti-PD-1 antibody nivolumab prolonged median overall survival in patients with chemotherapy-resistant pancreatic cancer compared to a lower dose of the same regimen. In this study, ATRA was the first drug to be found to cause ADAR1 degradation. We propose translation of a promising 2-pronged antitumor strategy using ATRA and nivolumab to convert immunologically “cold” into “hot” tumors susceptible to immune checkpoint blockade.

---

Interferon (IFN) signaling has pivotal roles in regulating antitumor immunity by directly activating immune responses and initiating programmed death of cancer cells (*1, 2*). The effects of IFNs are mediated through activation of JAK-STAT signaling pathways, followed by induction of IFN-stimulated genes (ISGs) (*3*). Simultaneously, IFNs also induce an early negative feedback loop that prevents overresponse of the immune system through upregulating inhibitory immune checkpoint pathways such as programmed cell death 1/programmed cell death ligand 1 (PD-1/PD-L1) and permitting cancer cells to escape immune surveillance (*4, 5*). Targeting PD-1/PD-L1 and other immune checkpoints has transformed the treatment of several cancers (*6*). However, low overall response rates and resistance in many tumors limit the universal use of checkpoint inhibitors in cancer treatment (*7*). Resistance to immune checkpoint inhibitors may occur because tumor cells evolve and create immunosuppressive microenvironments that are resistant to IFN signaling (*8, 9*). Although genomic defects in IFN pathway genes such as *IFNGR1* and *JAK1/2* in tumor cells have been reported to be a direct mechanism leading to immune checkpoint inhibitor resistance, wild-type IFN pathway genes have been found in some patients with resistant tumors (*8, 9*). These findings suggest that epigenetic underlying mechanisms are involved in dysregulating IFN pathways and allowing tumor cells to escape immune surveillance.

The ISG *ADAR* (double-stranded RNA-specific adenosine deaminase) encodes the protein ADAR1, an RNA-editing enzyme that limits the sensing of endogenous double-stranded RNA and negatively regulates IFN signaling. Recent data suggest that ADAR1 promotes immune suppression and enhances tumor viability in both enzymatic activity–dependent and –independent manners (*10, 11*). The orchestration of a tumor-promoting and immune-suppressive microenvironment by ADAR1 was supported by studies showing that tumor cells expressing high levels of ISGs, including *ADAR,* are sensitive to genetic suppression of ADAR1 (*12*) and that loss of ADAR1 expression in tumors increases their sensitivity to immune checkpoint inhibitors (*13, 14*). Thus, ADAR1 may act as an ideal target in tumors that display activation of IFN signaling. However, no specific inhibitor of ADAR1 is currently available, and it remains largely unknown how to antagonize ADAR1 in a way that can be translated into clinical use.

In this study, we first screened several common anticancer drugs for their ability to inhibit ADAR1. We found that all-trans retinoic acid (ATRA, also known as tretinoin) induced degradation of ADAR1 protein via ubiquitin-proteasome pathways in pancreatic and breast tumor cells. ATRA not only suppressed tumor growth by direct cytotoxic effects, but also provoked IFN signaling in tumor cells. Concomitantly, PD-L1, but not other immune checkpoints, was upregulated by ATRA through transcriptional activation in an RAR/RXR-dependent manner in pancreatic cancer cells. Moreover, combined treatment with ATRA and an anti-PD-1 therapeutic antibody improved survival in both pancreatic cancer and breast cancer mouse models, and repopulated infiltrating CD8+ T cells in the tumor microenvironment. More importantly, a higher dose of ATRA plus nivolumab prolonged median overall survival in patients with chemotherapy-resistant pancreatic tumors compared to patients treated with a lower dose of ATRA plus nivolumab. Thus, our study not only is the first to propose that ATRA can impede ADAR1 protein expression and upregulate PD-L1, but also highlights that ATRA plus nivolumab is a rational and promising therapeutic approach to treat cancer.

## Results

### ATRA reduced ADAR1 expression and stimulated immune response–related pathways in pancreatic cancer

Owing to the association between tumors’ ISG signature level and their ADAR1 dependence, we assessed expression profiles of ISGs (*13*) in various tumor cell types profiled in the Cancer Cell Line Encyclopedia (CCLE) to choose a tumor type with a high ISG signature for use as a model for screening of cancer drugs that might inhibit ADAR1 expression. The analysis revealed that upper aerodigestive tract tumors, pancreatic tumors, and urinary tract tumors had the 3 highest ISG z-scores (Figure S1A). The ADAR1 relative protein expression levels from CCLE are shown in Figure S1B. Pancreatic cancer was chosen as the model for further experiments because of the limited therapeutic choices and dismal prognosis of patients with advanced pancreatic cancer (*15*).

Considering that a drug repurposing approach may offer faster solutions than *de novo* drug design, we tested several common FDA-approved anticancer drugs for their ability to suppress ADAR1 expression: the chemotherapeutic agents 5-fluorouracil, cisplatin, and gemcitabine; the cyclin-dependent kinase (CDK) inhibitor dinaciclib; the receptor tyrosine kinase inhibitors erlotinib, sunitinib, and crizotinib; the Abl and Src family tyrosine kinase inhibitor dasatinib; the PARP inhibitor olaparib; the KRAS G12C inhibitor sotorasib; the differentiation agent ATRA; and compounds targeting inflammation-related molecules such as imiquimod (a TLR7 ligand) and tofacitinib, a JAK inhibitor. Among the initial 13 candidates, ATRA was the most significantly inhibited ADAR1 expression (Figure 1A, 1B, S2A and S2B). Accordingly, analysis of CCLE data also showed that the areas under the receiver operating characteristic curves for ATRA were inversely correlated with ADAR1 protein levels (R = −0.0989, *P* = 0.0366) (Figure S2C).

**Figure 1.**
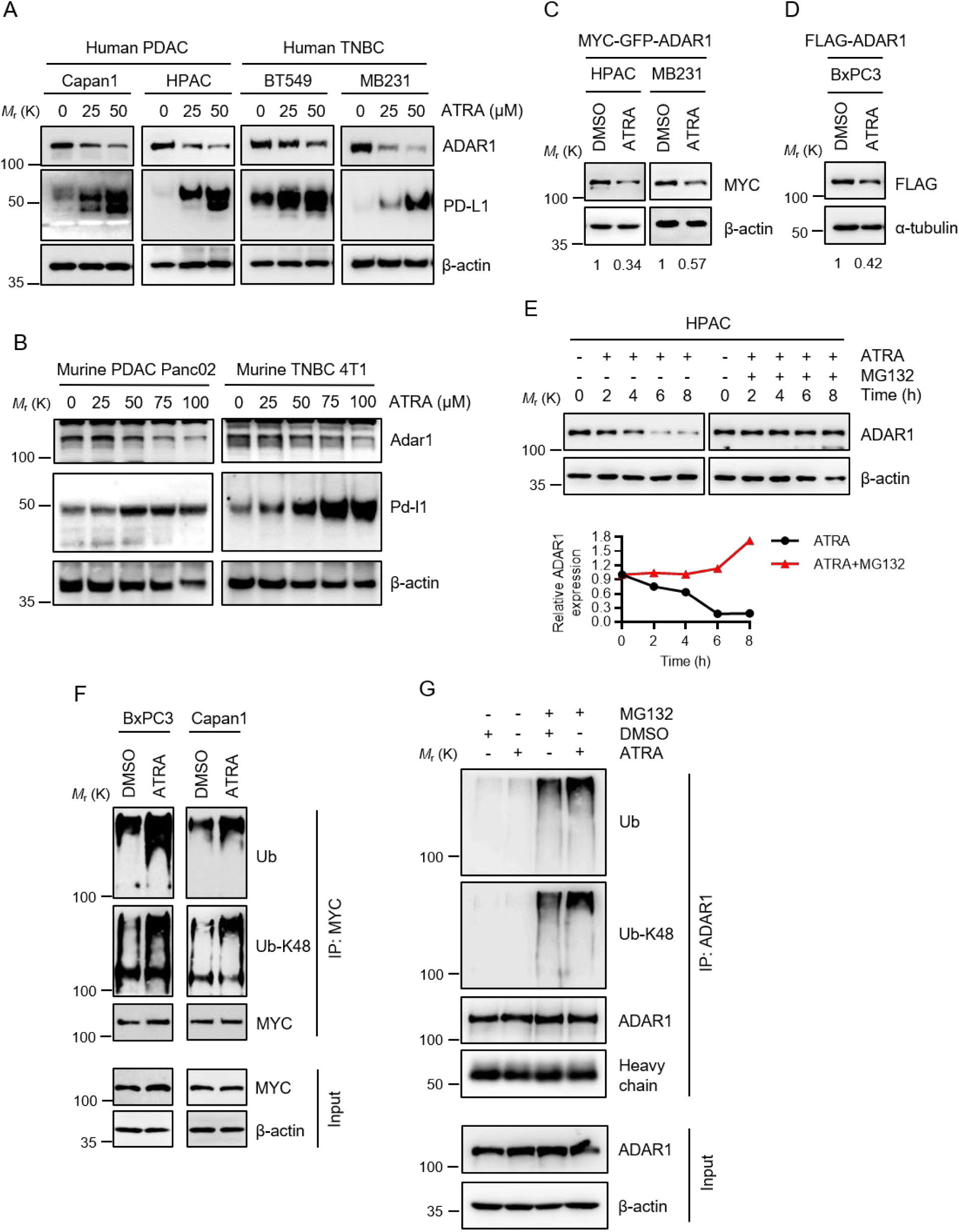
ATRA suppressed ADAR1 expression through the proteosome degradation pathway in pancreatic cancer cells. **(A)** Western blots showing the protein levels of ADAR1 and PD-L1 in Capan1, HPAC, BT549 and MB-231 cells treated with the indicated concentrations of ATRA for 24 h. PDAC, pancreatic ductal adenocarcinoma; TNBC, triple-negative breast cancer. **(B)** Western blots showing the protein levels of Adar1 and Pd-l1 in Panc02 and 4T1 cells treated with the indicated doses of ATRA for 24 h. **(C)** Western blots showing exogenous MYC-GFP-tagged ADAR1 protein expression in HPAC or MB-231 cells treated with ATRA for 24 h. **(D)** Western blots showing exogenous FLAG-tagged ADAR1 protein expression in BxPC3 cells treated with ATRA for 24 h. **(E)** Upper, Western blots showing the protein levels of ADAR1 in HPAC cells treated with ATRA or MG132 alone or in combination for the indicated time. Lower, Quantification of relative ADAR1 expression. **(F)** Co-immunoprecipitation (Co- IP) blots showing polyubiquitination (Ub) and K48-linked ubiquitination (Ub-K48) of ADAR1 in BxPC3 and Capan1 cells exogenously expressing MYC-GFP-tagged ADAR1 and treated with ATRA. **(G)** Co-IP blots showing polyubiquitination and Ub-K48 of ADAR1 in HPAC cells treated with ATRA or MG132.

Moreover, ATRA inhibited pancreatic cancer cell growth and colony formation (Figure S3A and S3B). Pancreatic cancer cell growth inhibited by ATRA was rescued by exogenous ADAR1 expression (Figure S3C and S3D), and knockdown of *ADAR* did not further sensitize pancreatic cancer cells to ATRA treatment (Figure S3E), suggesting that the reduction of pancreatic cancer cell viability induced by ATRA depended on loss of ADAR1.

Notably, ATRA stimulated activation of IFN-related downstream signaling molecules including STAT1, IRF3, and TBK1, as well as the immune checkpoint PD-L1 (Figure S2A and S2B). Analysis of data from the Kyoto Encyclopedia of Genes and Genomes (KEGG) showed that 2 of the positively enriched pathways mediated by ATRA were cytokine-cytokine receptor interaction and the JAK-STAT signaling pathway (Figure S4A and S4B). These findings were supported by analysis of the biological functions enriched by *ADAR* knockout (Figure S4C). Further, an analysis of disease-gene associations in the DisGeNET database indicated that ATRA is involved in a general response to inflammatory and infectious diseases (Figure S4D), suggesting that ATRA stimulates immune response–related pathways.

There are 2 ADAR1 isoforms, a constitutively expressed p110 isoform and an inflammatory cytokine response–induced p150 isoform (*16, 17*). Intriguingly, ADAR1-p110 expression was not affected by treatment with IFNs or ATRA (Figure S4E). However, unlike IFNs, ATRA significantly reduced ADAR1- p150 expression (Figure S4E).

PD-L1 has been reported to be upregulated by inflammatory stimuli such as IFN-β, IFN-γ and IL-6 (*18, 19*). We found that PD-L1 expression in pancreatic cancer and triple-negative breast cancer (TNBC; another immunologically cold tumor) cells was increased by treatment with IFNs or with ATRA (Figure 1A and 1B and Figure S4E). Unlike IFNs, which are known to cause increased expression of ISGs including *ADAR* that encodes ADAR1 which acts as a feedback brake on IFN response, ATRA stimulates a noncanonical immune response–related pathway, which in turn activated inflammatory response signaling molecules and PD-L1 while concomitantly reducing ADAR1.

### ATRA inhibited ADAR1 expression through ubiquitin-proteasome pathways

Next, we sought to demonstrate how ATRA inhibited ADAR1 expression. We found that ATRA did not reduce *ADAR* mRNA levels (Figure S5A); instead, it markedly decreased the expression of exogenous ADAR1 protein (Figure 1C and 1D), suggesting that ATRA regulates ADAR1 expression by posttranscriptional mechanisms.

The ubiquitin-proteasome and autophagy-lysosome systems are 2 major mechanisms involved in eukaryotic protein degradation. Treatment with the lysosome inhibitor chloroquine or the proteasome inhibitor MG132 showed that ADAR1 protein expression can be regulated by either lysosomal or proteasomal degradation (Figure S5B and S5C). However, in the pancreatic cancer cell lines BxPC3, HPAC and Capan1, MG132 mainly rescued ATRA-induced ADAR1 downregulation (Figure 1E and S5D and S5E). Co-immunoprecipitation showed that ATRA increased the level of ubiquitin and K48- linked polyubiquitination conjugation of ADAR1 in the presence of MG132 (Figure 1F and 1G). These results indicated that ubiquitin-proteasomal degradation is involved in ATRA-provoked ADAR1 degradation.

### ATRA induced PD-L1 transcriptional upregulation through RAR/RXR activation

Next, we addressed how ATRA induced PD-L1 expression. It has been well documented that ATRA acts as an agonist of 2 nuclear receptor families, retinoic acid receptor (RAR) and retinoid X receptor (RXR), which bind to DNA and directly regulate gene transcription (*20*). We found that ATRA upregulated *CD274* (PD-L1) mRNA levels in pancreatic cancer cells (Figure S6A), and that the transcription inhibitor actinomycin D reduced ATRA-induced PD-L1 protein levels (Figure S6B), suggesting that ATRA induces transcriptional upregulation of PD-L1. Consistently, exogenous PD-L1 expressed by a cDNA construct under a nonendogenous PD-L1 promoter was not increased by ATRA (Figure S6C), suggesting that ATRA does not upregulate PD-L1 via a posttranscriptional mechanism. AGN 193109, a high-affinity RAR antagonist, inhibited ATRA-induced PD-L1 expression (Figure S6D). Together, these results indicated that *CD274* (PD-L1) was transcriptionally upregulated by ATRA through RAR/RXR activation.

### Combination of ATRA and anti-PD-1 antibody improved antitumor effect in a murine pancreatic cancer model

Because ATRA inhibits ADAR1 and induces PD-L1, we reasoned that combining these 2 targets might improve therapeutic efficacy. To explore opportunities for improving therapies for pancreatic cancer by combining ATRA and an anti-PD-1 antibody, we first tested whether ATRA stimulated functional PD-L1 expression. We found that cell surface PD-L1 expression increased under ATRA treatment (Figure S6E). Immune checkpoint arrays showed that apart from PD-L1, no other major immune checkpoint molecules were obviously regulated by ATRA (Figure S6F). These results suggested that combined treatment with ATRA and anti-PD-1 blockade may be a rational therapeutic approach to pancreatic cancer.

Because ATRA inhibited pancreatic cancer growth through downregulating ADAR1 expression and increasing PD-L1 expression, we further tested whether ATRA and anti-PD-1 blockade would synergize in pancreatic cancer models. To this end, we analyzed the *in vivo* efficacy of combination therapy with ATRA and an anti-PD-1 antibody in the murine syngeneic Panc02 pancreatic cancer model (Figure 2A). Mice treated with ATRA plus the anti-PD-1 antibody had smaller tumors and longer survival time than did mice treated with either agent as monotherapy (Figure 2B, 2C, 2D and 2E).

**Figure 2.**
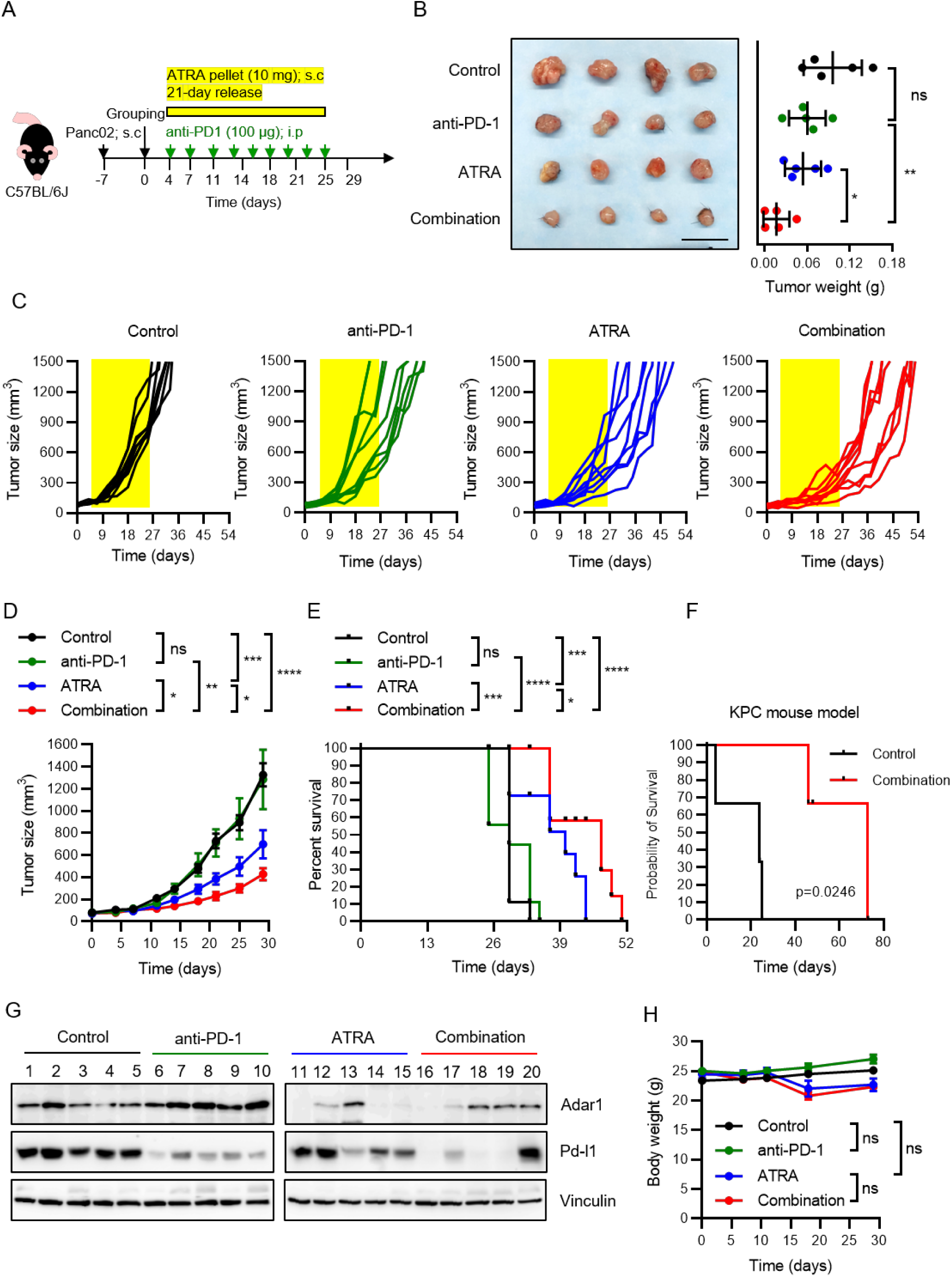
Combined treatment with ATRA and an anti-PD-1 antibody enhanced antitumor effect in a murine pancreatic cancer model. **(A)** Schematic representation of the treatment schedule in the Panc02 mouse tumor model. Panc02 tumor-bearing mice were given ATRA or placebo pellets (10 mg per mouse, subcutaneously [s.c.], 21-day release). Administration of anti-PD-1 antibody or IgG control (100 μg per mouse, intraperitoneally [i.p.], 3 times weekly) was started along with ATRA or placebo pellets. Tumor growth, body weight, and mouse survival were measured. **(B)** Left, representative images showing Panc02 tumors harvested from mice that received the indicated treatments. Scale bar, 1.2 cm. Right, tumor weight. Data represent mean ± SD. **(C)** Tumor growth curves of individual mice injected with Panc02 tumor cells and subjected to the indicated treatments. **(D)** Mean tumor growth in mice injected with Panc02 tumor cells and subjected to the indicated treatments. Each dot represents the mean tumor size from 9 mice in each treatment group; whiskers represent SD. **(E)** Survival curves of mice that received the indicated treatments (9 mice/group). **(F)** Survival curves of KPC mice that received the indicated treatments (3 mice/group). **(G)** Western blots showing Adar1 and Pd-l1 protein expression levels in Panc02 cells from mice treated with the indicated regimens. **(H)** Body weight of tumor-bearing mice treated with the indicated treatments. Data are shown as mean ± SD. \**P* < 0.05; \*\**P* < 0.01; \*\*\**P* < 0.001; \*\*\*\**P* < 0.0001; ns, not significant.

We used genetically engineered KPC mice, a model that recapitulates human pancreatic cancer, to characterize disease progression. Our findings indicated that the combination of ATRA and anti-PD-1 antibody significantly (*P* = 0.0246) improved mouse survival (Figure 2F). Moreover, inhibition of ADAR1 expression by ATRA was confirmed in a syngeneic Panc02 mouse model (Figure 2G). The average mouse body weight, kidney function, and liver function did not significantly differ among the 4 experimental groups (Figure 2H and S7D). To confirm the effectiveness of the combination strategy, we performed similar experiments in a 4T1 orthotopic mouse mammary cancer model and obtained similar results (Figure S7A, S7B and S7C). Altogether, these results suggested that combined treatment with ATRA and an anti-PD-1 antibody could be a promising therapeutic approach to cancer therapy without obvious toxic effects.

To identify alterations of the immune-cell profiles of the tumor microenvironments under different treatment scenarios, we used time-of-flight mass cytometry (CyTOF). Unsupervised clustering analysis of CD45+ immune cells from tumors using viSNE and FlowSOM identified 9 major immune-cell populations (Figure 3A, 3B, 3C, S8A, S8B, S8C and S8D). Among these populations, the CD8+ T-cell population was visually distinct across all treatment groups (Figure 3C and 3D).

**Figure 3.**
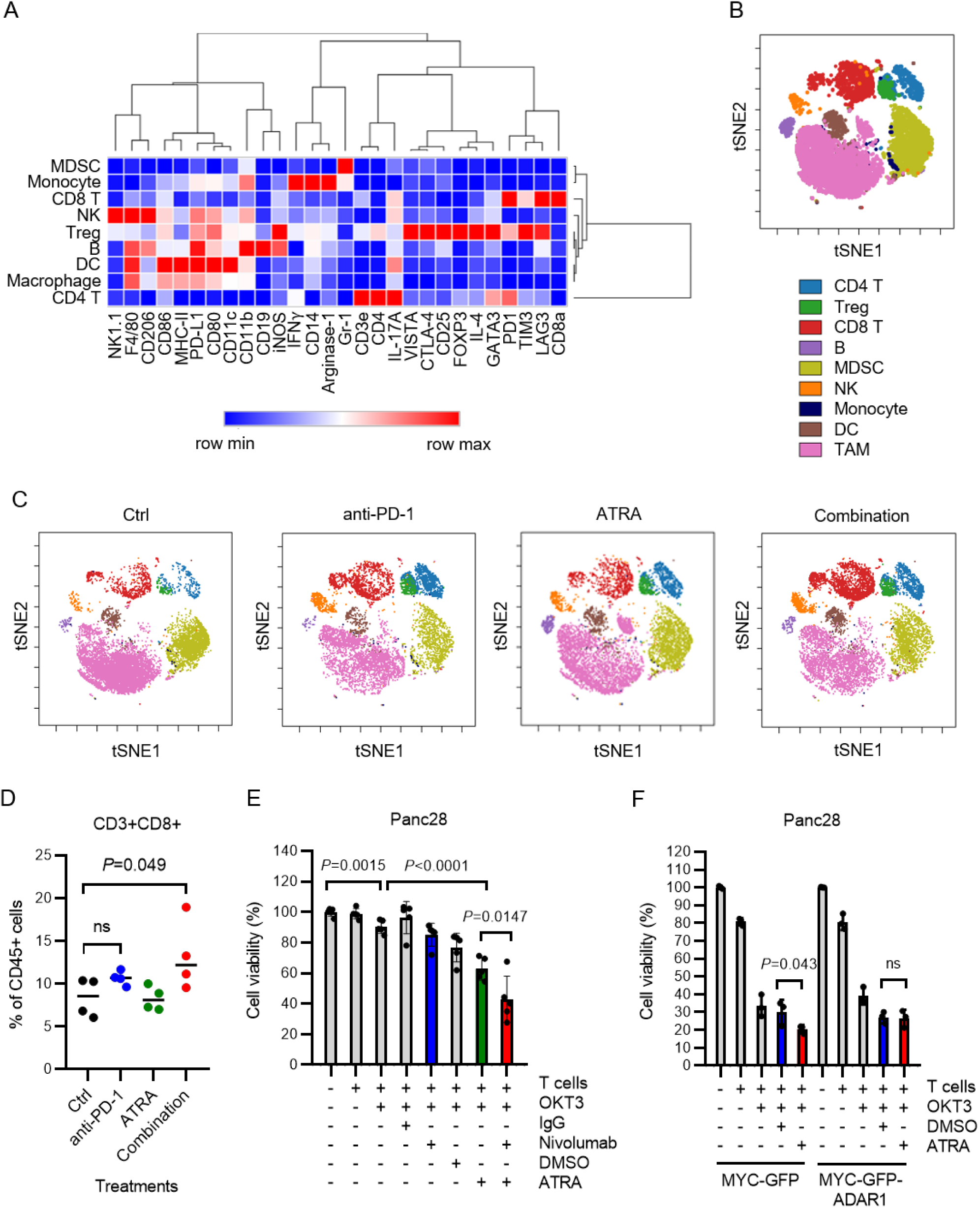
Combination of anti-PD-1 and ATRA targets specific subsets of tumor-infiltrating immune cells. **(A)** Heatmap showing differential marker expression in CD45+ tumor-infiltrating lymphocyte (TIL) clusters identified by analysis of time-of-flight mass cytometry data with viSNE and FlowSOM. DC, dendritic cell; MDSC, myeloid-derived suppressor cell; Treg, regulatory T cell; NK, natural killer cell. **(B)** Annotation of TIL populations based on differential marker expression as shown in the heatmap (A) and Figure S8. TAM, tumor-associated macrophage. **(C)** Cellular distribution and clustering as defined by tSNE1 and tSNE2, colored by cell phenotype, for Panc02 tumors subjected to the indicated treatments. Data show all normalized viable single cells; analysis was conducted using viSNE and FlowSOM. **(D)** Percentage of CD8+ T cells among total CD45+ TILs under the indicated treatments. **(E)** T-cell cytotoxicity (CCK-8) showing effects of the indicated treatments on pancreatic cancer cell viability. Target Panc28 cancer cells were engineered to express Fcγ receptor 2A fused with the luciferase Luc2 at the C-terminus (Panc28-RL2), and then incubated with CD8+ T cells and the CD3 antibody OKT3 for 3 days. Data represent mean ± SD. *P* values were calculated by Student *t* test. **(F)** T-cell cytotoxicity (CCK-8) assay showing effects of ATRA on pancreatic cancer cell viability. Panc28-RL2 cells overexpressing MYC-GFP-ADAR1 or MYC-GFP (control) were incubated with CD8+ T cells and the CD3 antibody OKT3 for 3 days. Data represent mean ± SD. *P* values calculated by Student *t* test.

To validate that CD8+ T cells contributed to the observed therapeutic effects, we used a previously established *in vitro* co-culture system to assess engineered CD8+ T-cell cytotoxicity toward tumor cells (*21*). Panc28 human pancreatic cancer cells were transduced to express the extracellular domain of FcγR2a and luciferase (Figure S8E). The addition of the anti-CD3 antibody OKT3 was expected to trigger CD8+ T-cell killing of Panc28 cells by simultaneously engaging CD8+ T cells and tumor cells. Using this system, we showed that ATRA promoted tumor-cell killing mediated by CD8+ T cells, and that the combination of ATRA and the FDA-approved PD-1 immune checkpoint inhibitor nivolumab further enhanced the tumor cell–killing efficacy of CD8+ T cells (Figure 3E).

Because ATRA inhibited ADAR1 expression (Figure 1A, 1B, S2A and S2B), we established that Panc28 cells overexpressed ADAR1 to investigate whether ADAR1 negatively contributed to ATRA’s promotion of CD8+ T-cell cytotoxicity (Figure S8F). Compared to the vehicle control group, Panc28 cells overexpressing a MYC-GFP control vector exhibited more CD8+ T-cell killing activity in the presence of ATRA. However, this CD8+ T-cell killing activity was compromised when ADAR1 was overexpressed (Figure 3F). These results revealed that ATRA-mediated downregulation of ADAR1 is required for ATRA- mediated CD8+ T-cell cytotoxicity.

Notably, combined treatment with ATRA and an anti-PD-1 antibody *in vivo* concomitantly stimulated recruitment of immune cells with antitumor function other than CD8+ T cells, for example T helper 1 (Th1) cells and CD80+ M1 macrophages (Figure S8G and S8K). On the other hand, the combined treatment inhibited or did not affect immunosuppressive cells such as regulatory T cells, M2 macrophages, and myeloid-derived suppressor cells (Figure S8I, S8L and S8M). In addition, significant increases of immune checkpoint molecules, including CTLA-4, LAG3, TIM-3, VISTA, PD-1, and PD-L1 expressed on tumor-infiltrating lymphocytes (TILs), were detected in the combined treatment group (Figure S9), indicating that the combined treatment provoked a T cell-inflamed tumor microenvironment. Thus, the data revealed that the combination of ATRA and an anti-PD-1 antibody may reprogram the tumor microenvironment to switch certain pancreatic cancers from “cold” into “hot” tumors via recalling antitumor immunity.

### Combination of ATRA and nivolumab lengthened median overall survival in patients with chemotherapy-refractory pancreatic cancer

The encouraging *in vivo* results (Figure 2, 3, S7, S8 and S9) prompted us to investigate the clinical relevance of ADAR1. Analysis of data from The Cancer Genome Atlas (TCGA) showed that *ADAR* mRNA was upregulated in various cancers including pancreatic cancer and breast cancer (Figure 4A). High expression of ADAR1 protein was associated with poor survival both in patients with pancreatic cancer (*P* = 0.0045) (Figure 4B and 4C) and those with breast cancer (*P* = 0.0162 and *P* = 0.0373 in 2 independent cohorts) (Figure 4D).

**Figure 4.**
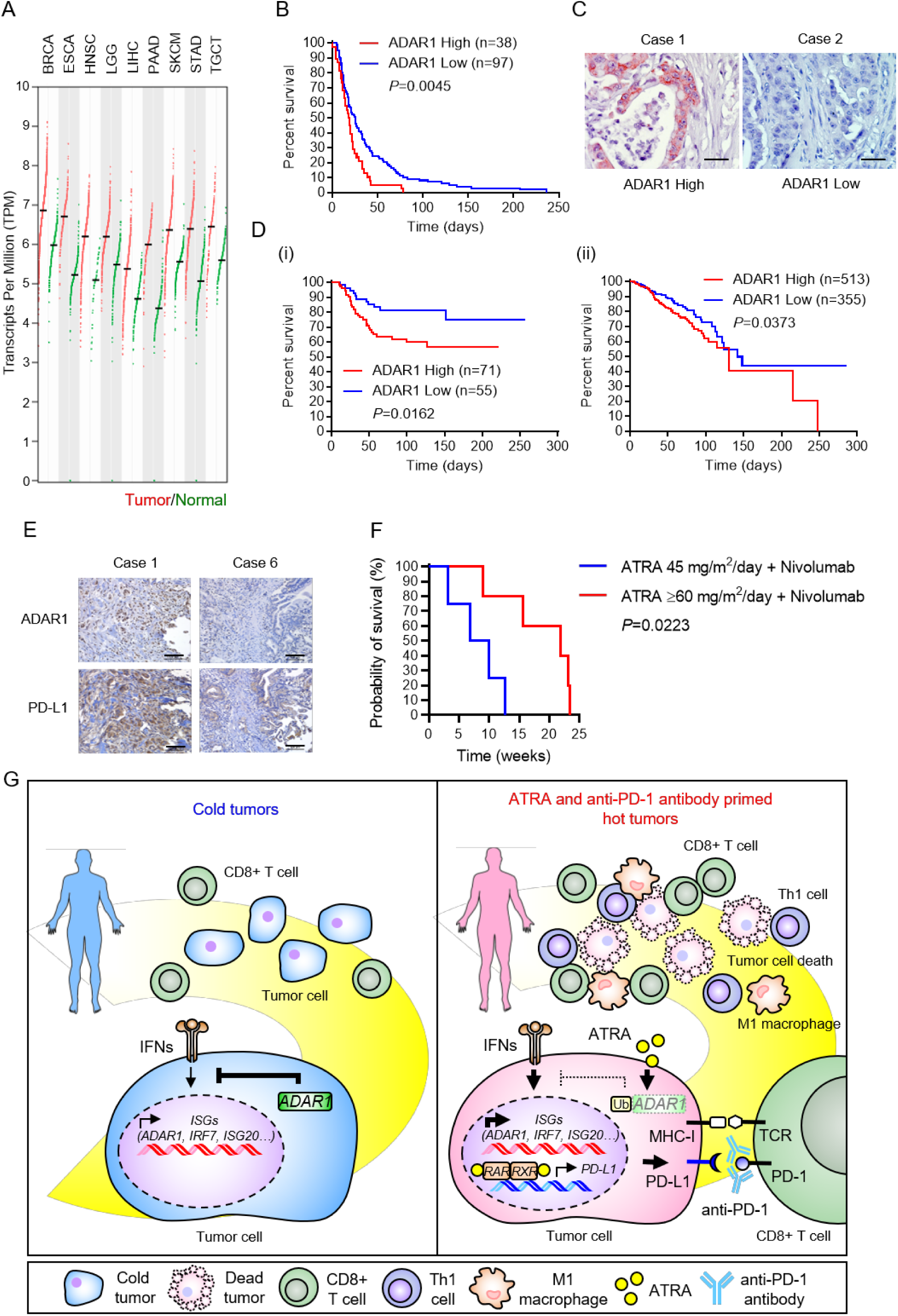
ADAR1 expression positively correlates with poor survival in patients with cancer. **(A)** *ADAR* gene expression profiling by GEPIA2. Red dots: tumor tissues; green dots: normal tissues. The bars represent the median expression of certain tumor type or normal tissue. BRCA, invasive breast carcinoma; ESCA, esophageal carcinoma; HNSC, head and neck squamous cell carcinoma; LGG, lower-grade brain glioma; LIHC, liver hepatocellular carcinoma; PAAD, pancreatic adenocarcinoma; SKCM, skin cutaneous melanoma; STAD, stomach adenocarcinoma; TGCT, testicular germ cell tumor. **(B)** Kaplan-Meier overall survival curves of 135 pancreatic cancer patients in the cohort with high and low ADAR1 expression determined by immunohistochemical staining. **(C)** Representative images of the immunohistochemical staining of ADAR1 in tumor tissue from patients in the MDA cohort. Scale bars, 50 μm. **(D)** Survival curves of breast cancer patients from 2 independent cohorts by ADAR1 expression level. (i) Liu *et al.*, 2014; (ii) TCGA. **(E)** Representative images of the immunohistochemical staining of ADAR1 and PD-L1 in tumor tissue from pancreatic cancer patients before treatment with ATRA and nivolumab. Scale bars, 100 μm. **(F)** Kaplan-Meier overall survival curves for 9 pancreatic cancer patients treated with ATRA 45 mg/m^2^/day plus nivolumab or ATRA ≥60 mg/m^2^/day plus nivolumab. **(G)** Schematic illustration of the mechanism of switching “cold” (immune-excluded) tumors to “hot” (immune-infiltrated) tumors by using ATRA and anti-PD-1 antibody.

The above encouraging preclinical data from *in vitro* systems and multiple animal models prompted us to initiate a pilot clinical trial to evaluate the antitumor activity of ATRA plus nivolumab in patients with advanced pancreatic cancer whose tumors had progressed during chemotherapy. Since 2021, 9 patients (4 male and 5 female) have participated in the trial (Table S1). The participants’ median age was 63 years (range, 54-79). All patients had metastatic pancreatic adenocarcinoma that had progressed after treatment with a median of 3 (range, 2-4) lines of chemotherapy, including a gemcitabine-containing regimen.

Immunochemistry staining was used to determine ADAR1 and PD-L1 expression before the experimental treatments (Figure 4E). Patients received from 1 to 11 2-week cycles (median of 4) of ATRA plus nivolumab. 5 patients received escalated doses of ATRA if the initial dose of 45 mg/m^2^/day was well tolerated: 60 mg/m^2^/day in 2 patients and 75 mg/m^2^/day in 3 patients. The best response was stable disease in 3 (33%) patients (Table S1). The median overall survival time was 21.9 weeks (95% CI, 8.4-35.4) for patients receiving a higher dose of ATRA (≥60 mg/m^2^/day), compared to 6.9 weeks (95% CI, 0.1-13.6) for patients receiving ATRA at a dose of 45 mg/m^2^/day (*P* = 0.0223; Figure 4F). Thus, the clinical results suggested that a combination of a higher dose (≥60 mg/m^2^/day) of ATRA with nivolumab improved the overall survival time in patients with advanced pancreatic cancer. A model depicting the proposed mechanisms of this therapeutic effect is shown in Figure 4G, which illustrates how ATRA and an anti-PD- 1 antibody could convert “cold” to “hot” tumors and attract cytotoxic CD8+ T cells to kill tumor cells.

### Increased levels of ADAR1 and ISGs were observed in gemcitabine-resistant pancreatic cancer cells

Gemcitabine is one of the fundamental first-line drugs for pancreatic cancer treatment; however, a substantial portion of pancreatic cancers are resistant to gemcitabine (*22*). We therefore asked whether the newly identified mechanisms might contribute to gemcitabine resistance. We found that gemcitabine induced expression of PD-L1 *in vitro*, and the addition of ATRA both inhibited ADAR1 and further induced PD-L1 in gemcitabine-treated pancreatic cancer cells (Figure S10A). Thus, to examine whether ATRA could overcome gemcitabine resistance, we developed gemcitabine-resistant Panc28 and Capan1 pancreatic cancer cell lines. Intriguingly, ATRA repressed the viability of gemcitabine-resistant cells (Figure S10B). Of note, gemcitabine-resistant Panc28 and Capan1 cells had higher or unchanged ADAR1 expression levels and lower PD-L1 expression levels compared to their corresponding parental cells (Figure S10C). Although gemcitabine did not apparently affect ADAR1 and PD-L1 expression in resistant cells, ATRA significantly inhibited ADAR1 and induced PD-L1 (Figure S10C). Moreover, the results of CD8+ T-cell cytotoxicity assays revealed that combined treatment with ATRA and nivolumab augmented CD8+ T-cell cytotoxicity against gemcitabine-resistant cells (Figure S8E and red line in Figure S10D).

Next, we conducted Gene Set Enrichment Analysis to investigate the gene sets associated with gemcitabine-resistant pancreatic cancer. The results indicated that gemcitabine-resistant pancreatic cancers have positively enriched immune-related gene sets, including IFN-α, IFN-γ, inflammatory response, TNF-α signaling via NFκB, and IL6-JAK-STAT3 signaling (normalized enrichment scores = 2.452, 2.251, 2.076, 1.993, and 1.758, respectively) (Figure S11A; GSE79953). The heatmaps of ISGs from 2 independent NCBI GEO databases (GSE79953 and GSE140077) indicated that ISGs including *ADAR* were highly enriched in gemcitabine-resistant pancreatic cancer cells (Figure S11B and S11C). Analysis of TCGA databases also showed gemcitabine resistance gene signature positively correlated with ISG gene signature and *ADAR* in pancreatic, breast, lung and bladder cancer (Figure S11D and S11E). These findings indicated that although IFN responses, which contribute to cytotoxic and antitumor functions, were stimulated in gemcitabine-resistant cells, ADAR1, which plays an essential role in promoting tumor-cell escape from antitumor immunity, was also upregulated in gemcitabine-resistant cells (Figure S10C, S11A S11B and S11C). Thus, a combination of ATRA and an anti-PD-1 antibody may be a promising therapeutic strategy for pancreatic cancer patients in whom gemcitabine chemotherapy has failed.

Taken together, our findings demonstrated that pancreatic cancer highly expresses ISGs including *ADAR* which encodes ADAR1 protein. ATRA significantly inhibited ADAR1 expression through ubiquitin-proteasome pathways and concomitantly induced PD-L1 gene expression through RAR/RXR heterodimers. The combination of ATRA and an anti-PD-1 antibody stimulated CD8+ T-cell infiltration in the tumor microenvironment and impeded tumorigenicity in both pancreatic cancer and breast cancer mouse models (Figure 4G). The results of our pilot clinical study supported those of our preclinical models, demonstrating the promise of this therapeutic approach for chemotherapy-resistant pancreatic cancer.

## Discussion

Our data provide a proof of concept for the combination therapy to treat pancreatic cancer by targeting ADAR1 and PD-L1, which reinvigorated anti-tumor response through converting immunologically “cold” into “hot” tumor. ADAR1 plays a pivotal role in maintaining self-tolerance and preventing autoimmunity (*23, 24*). On one hand, it has recently been reported that ADAR1 contributes to tumorigenesis through suppressing IFN response and innate immunity by endogenous RNA editing (*12–14*). On the other hand, ADAR1 can promote tumorigenesis by causing genetic and/or epigenetic alterations via editing certain coding regions of mRNA, splicing branch sites and microRNAs, processing Z-RNA through enzyme-dependent mechanisms, or directly interacting with Dicers to promote processing of small interfering RNAs and microRNAs through enzyme-independent pathways (*25*). Accumulated evidence indicates that intratumoral ADAR1 inhibition or depletion can be a promising strategy for cancer treatment (*12–14*). Despite reports that SUMO-1 and Lys-48 ubiquitination modifications contribute to impaired enzymatic activity of adenosine deaminase and protein degradation of ADAR1, respectively (*26, 27*), there is limited understanding of ADAR1 protein modifications and degradation and a lack of translational studies.

ATRA has been reported to play important roles in cell development and differentiation, immune regulation, and tumor progression (*28–30*). Moreover, ATRA has been shown to reprogram pancreatic stellate cells and ameliorate pancreatic fibrosis (*31*). Currently, ATRA is commonly used for the treatment of acute promyelocytic leukemia (*32*), but it does not show significant efficacy in solid tumors (*33, 34*). This may be because PD-L1 is also induced by ATRA; therefore, using an anti-PD-1 antibody in combination with ATRA may improve ATRA-based treatment of solid tumors by overcoming their immunosuppressive microenvironment.

In this study, we found that ATRA significantly reduced ADAR1 expression through ubiquitin-proteasome pathways and concomitantly stimulated the IFN-related response in tumor cells. Although ATRA, like IFNs, increased PD-L1 expression at transcription levels, compromising antitumor immunity, this increase in PD-L1 expression can be overcome by anti-PD-1 antibodies. Vice versa, the elevated ADAR1 expression triggered by the enhanced IFN response upon PD-1/PD-L1 blockade can be suppressed by ATRA. Hence, a combination of ATRA and PD-1/PD-L1 inhibitors could be a rational strategy to improve both drugs’ efficacies against cancer. Additionally, the sustained IFN response produced by antagonizing both PD-1/PD-L1- and ADAR1-mediated pathways could minimize tumor resistance or relapse. Indeed, our pancreatic cancer and breast cancer syngeneic mouse models, and KPC mouse model indicated that combined treatment with ATRA and an anti-PD-1 antibody synergistically suppressed tumor growth and prolonged survival. In the pancreatic cancer model, the combination of ATRA and anti-PD-1 blockade increased the number of infiltrating CD8+ T cells, Th1 cells, and M1-like macrophages in the tumor microenvironment. Numerous immune checkpoints were highly expressed on TILs under treatment conditions, suggesting that ATRA plus an anti-PD-1 antibody may turn nonimmunogenic “cold” tumors into immunogenic “hot” ones.

The dismal prognosis of pancreatic cancer patients in whom chemotherapy fails is reflected by the short median post-progression survival durations of 1.3 to 3.0 months in trials of second-line chemotherapy (*35–37*). Novel immunotherapy approaches, including PD-1/PD-L1 blockade to provoke antitumor immunity, have failed in treating pancreatic cancer patients (*38*). In our pilot clinical trial of concurrent targeting of ADAR1 and PD-1 in patients with advanced, chemotherapy-refractory pancreatic cancer, patients treated with a high dose of ATRA plus nivolumab had significantly longer median overall survival (21.9 weeks) than did those treated with a low dose of ATRA plus nivolumab (6.9 weeks). In addition, we found that ADAR1 expression was increased in gemcitabine-resistant pancreatic cancer cells and was repressed by ATRA. CD8+ T-cell killing assays indicated that ATRA alone promoted cytotoxic activity toward gemcitabine-resistant cells and that adding nivolumab further enhanced the effect. Taken together, the results of our preclinical and clinical studies suggest that targeting both ADAR1 and PD-1/PD-L1 may restimulate an IFN response to improve the outcomes of immunotherapy for patients with gemcitabine-resistant pancreatic cancer.

In addition to regulating gene expression via RAR/RXR heterodimers at the transcriptional level, a small number of reports indicate that ATRA regulates protein expression through ubiquitin-proteasome pathways and regulation of kinase activity (*39*). Future studies are needed to investigate the underlying mechanism by which ATRA suppresses ADAR1 expression, whether through activating an unknown E3 ubiquitin ligase or inactivating deubiquitinases.

In summary, we discovered 3 novel antitumor mechanisms of ATRA: inhibiting ADAR1, upregulating PD-1 expression, and enhancing IFN signaling in tumor cells. The combinatorial approach targeting ADAR1 and PD-L1 using ATRA and nivolumab, which reinvigorated the antitumor response, showed promise in treating patients with chemotherapy-refractory pancreatic cancer. For translation of an ATRA- plus-nivolumab regimen to the clinic, the regimen will require further investigation and validation in clinical trials with a larger population of patients and different tumor types.

## Materials and Methods

### Cell lines and plasmids

4 human pancreatic cancer cell lines (BxPC3, Capan1, HPAC, and Panc28), 2 TNBC cell lines (BT-549 and MDA-MB-231), a human embryonic kidney cell line (HEK293T), a murine pancreatic cancer cell line (Panc02), and a murine mammary cancer cell line (4T1) were used in this study. The Capan1 and Panc28 cells which were resistant to gemcitabine were generated by selection under increasing gradient of gemcitabine. The gemcitabine resistant cells were than cultured in periodically added 1 μM gemcitabine to maintain cell resistance. The pCDH-CMV-MYC-ADAR1-Neo expression vector was generated by inserting the full-length ADAR cDNA (NM_001111) purchased from Addgene into the pCDH-CMV-MCS-EF1α-Neo vector obtained from Addgene. The pMA2642-FCGR2A-Luc2 expression vector was used for T-cell killing assays. The pLKO.1-shLuc control plasmid (TRCN0000072243), pLKO.1-shLacZ control plasmid (TRCN0000072237), and *ADAR* short hairpin RNAs (TRCN0000301036 and TRCN0000050788), used for gene silencing, were obtained from Merck Millipore.

### Reagents

5-Fluorouracil, AGN 193109, dasatinib, olaparib, sotorasib, tofacitinib, MG132, and Cell Counting Kit-8 (CCK-8) were purchased from MedChemExpress. Cisplatin, crizotinib, dinaciclib, erlotinib, gemcitabine, and sunitinib were purchased from Selleck Chemicals. Chloroquine and all-trans retinoic acid (ATRA, tretinoin) were purchased form Millipore Sigma. Actinomycin D and imiquimod were purchased from Cayman. Recombinant human IL-2 (carrier-free) was purchased from BioLegend. Human Immune Checkpoint Array C1 was purchased from RayBiotech. The iTaq Universal SYBR Green Supermix and the iScript Reverse Transcription Supermix for reverse transcription–real-time PCR (RT- qPCR) were purchased from Bio-Rad. Matrigel Growth Factor Reduced Basement Membrane Matrix was purchased from Corning. ATRA and placebo pellets (10 mg/pellet; 21-day release) were purchased from Innovative Research of America.

### Antibodies

APC mouse IgG2b, κ isotype control antibody, APC anti-human CD274 (B7-H1, PD-L1) antibody, PE mouse IgG1, κ isotype control (FC) antibody, PE anti-human FCGR2A/CD32A antibody, purified anti-human CD3 antibody (OKT3), and anti-human CD28 antibody were purchased from BioLegend. Human ADAR1 (p150 isoform) antibody was purchased from Bethyl Laboratories, Inc. Human ADAR1 (p110 isoform) antibody and vinculin antibody were purchased from Proteintech. Mouse Adar1 antibody, IRF3 antibody and phospho-IRF3-S386 antibody were purchased from ABclonal. Mouse Pd-l1 antibody was purchased from R&D Systems. α-Tubulin antibody, β-actin antibody for immune blotting assays, and monoclonal ANTI-FLAG M2 antibody for immunoprecipitation assays were purchased from Millipore Sigma. Anti-ubiquitin antibody (P4D1) was purchased from Santa Cruz Biotechnology. Myc-Tag (71D10) rabbit monoclonal antibody, K48-linkage specific polyubiquitin (D9D5) rabbit monoclonal antibody, human PD-L1 (E1L3N) XP rabbit monoclonal antibody, PARP (46D11) rabbit monoclonal antibody, Stat1 (D1K9Y) rabbit monoclonal antibody, phospho-Stat1 (Ser727) antibody, TBK1/NAK (D1B4) rabbit monoclonal antibody, and phospho-TBK1/NAK (Ser172) (D52C2) XP rabbit monoclonal antibody were purchased from Cell Signaling Technology. LC3B antibody, ADAR1 antibody and PD-L1 antibody were purchased from Abcam. Human IFN-β and human IFN-γ recombinant proteins were purchased from Sino Biological. InVivoMAb anti-mouse PD-1 (CD279) and InVivoMAb rat IgG2a isotype control antibodies for animal experiments were purchased from Bio X Cell.

### Lentiviral transduction

The psPAX2, pMD2.G, and lentiviral expression vectors (pCDH-Neomycin, pCDH-MYC-EGFP-ADAR1, and pCDH-FLAG-ADAR1 clones) were co-transfected into HEK293T cells overnight to package the virus. After the medium was replaced, the virus-containing supernatant was collected at day 2 after transfection. For virus transduction, the cells were mixed with virus-containing supernatants supplemented with 8 μg/mL polybrene (Thermo Fischer Scientific). After infection with the virus for 24 h, the infected cells were selected with 1 mg/mL neomycin for 3 days.

### Animal experiments

C57BL/6 mice and BALB/cJ mice we used in this study were purchased from the Jackson Laboratory. P48Cre+ LKRAS+ P53L/+ (KPC) mice were gifts from Dr. Haoqiang Ying. For the pancreatic cancer mouse model, 1 × 105 Panc02 cells were subcutaneously implanted on the backs of syngeneic C57BL/6 mice. For the breast cancer mouse model, 1 × 105 4T1 cells were orthotopically implanted into the mammary fat pads of syngeneic BALB/cJ mice. Mice were randomly divided into groups when treatment started 10 days after tumor implantation. ATRA or placebo pellets with reasonable dosage (10 mg/pellet; 21-day release) were subcutaneously implanted into the dorsal necks of the mice. For administration of anti-PD-1 antibody, the mice were treated 3 times per week for 21 days with intraperitoneal injection of either 100 μg anti-PD-1 antibody in phosphate-buffered saline (PBS) or its corresponding IgG in PBS as control. Each group contained 5-9 mice. Tumor size was measured using calipers and calculated using the following formula: tumor volume = L × W2/2 (L, length; W, width). The body weight of each mouse was measured at least twice per week. For survival analyses, tumor volumes greater than 1000 mm3 for the pancreatic cancer mouse model and 700 mm3 for the breast cancer mouse model were chosen as endpoints. For the P48Cre+ LKRAS+ P53L/+ (KPC) mouse model, ATRA or placebo pellets were given after pancreatic cancer formation was confirmed by magnetic resonance imaging. Each group contained 3 mice. When KPC mice became moribund from cancer, they were euthanized, and euthanasia served as the death endpoint for survival analysis. For blood chemistry analysis and CyTOF analysis, blood and tumors, respectively, were collected from mice bearing Panc02 tumors after the completion of treatment. The Department of Veterinary Medicine and Surgery at The University of Texas MD Anderson Cancer Center performed blood chemistry analysis using its standard operating procedures. All study protocols were approved by the Institutional Animal Care and Use Committee at The University of Texas MD Anderson Cancer Center.

### Immunophenotyping of tumor-infiltrating leukocytes by CyTOF

CyTOF was carried out as previously described (*21*). In brief, fresh Panc02 tumors were collected from mice bearing syngeneic subcutaneous tumors after treatment completion. Tumor samples were dissociated using the GentleMACS system (Miltenyi Biotec) according to the manufacturer’s protocols. Dissociated cells were stained with Cell-ID cisplatin (Fluidigm) for viability analyses. Cells were then stained with heavy metal isotope-conjugated antibodies to immune cell surface proteins (Table S2). For intracellular staining, cells were fixed and permeabilized with eBioscience Foxp3/Transcription Factor Staining Buffer Set (Thermo Fisher Scientific) and then incubated with heavy metal isotope-conjugated antibodies (Table S2). Cell-ID Intercalator-Ir (Fluidigm) was used to distinguish live and single-nucleated cells. For CyTOF data acquisition, cells were fixed with 1.6% paraformaldehyde, pelleted in water, and then analyzed on a Helios mass cytometer (Fluidigm) at the MD Anderson Flow Cytometry and Cellular Imaging Core Facility. Mass cytometry data files were analyzed using Cytobank analysis software (https://www.cytobank.org/). Markers of TILs were gated and clustered with the viSNE algorithm. The population of TILs was subsequently clustered using the FlowSOM algorithm. The markers for identifying major TIL subsets are shown in Table S3.

### Flow cytometric analysis

For antibody staining of PD-L1 or FcγR2a cell surface proteins, cells were trypsinized and incubated with fluorescence-labeled antibodies in 50 μL 1% bovine serum albumin (BSA) in PBS staining/wash buffer for 30 min in the dark. Cells were then washed and resuspended in 0.5 mL staining/wash buffer, and flow cytometry was performed. Data were acquired using a BD FACSCanto II flow cytometer with BD FACSDiva 8.0.1 software (BD Biosciences) and analyzed with FlowJo software (Becton Dickinson).

### Immune checkpoint array

Detection of immune checkpoints present in ATRA-treated HPAC cells was done using antibody array-based technology (Human Immune Checkpoint Array C1, RayBiotech, Inc.). Briefly, HPAC cells treated with or without 50 μM ATRA for 24 h were lysed using cell lysis buffer. Blocking, protein sample incubation, biotinylated antibody cocktail incubation and HRP-streptavidin incubation were performed following the manufacturer’s instructions. Images were visualized using the ImageQuant LAS 4000 imaging system (GE). ImageJ software was used for densitometry measurements of the signal.

### RT-qPCR

From each sample, 1 μg total RNA was retrotranscribed using iScript Reverse Transcription Supermix (Bio-Rad) based on the manufacturer’s instructions. All the primer sequences are shown in Table S4. iQ SYBR Green Supermix (Bio-Rad) was used for 20-μL qPCR reactions. Each 10-μL reaction contained 10 μL iQ SYBR Green Supermix, 0.5 μM of forward and reverse primers, and 10 μL of 20-fold diluted cDNA. The qPCR was performed on the Applied Biosystems Real-Time PCR Instrument (Thermo Fisher Scientific).

### T cell–mediated cytotoxicity assay

T cell–mediated cytotoxicity assays were carried out as previously described (*21*). CD8+ T cells were isolated from human peripheral blood mononuclear cells (STEMCELL Technologies) using the EasySep Human CD8+ T Cell Isolation Kit (STEMCELL Technologies). Target cells Panc28, Panc28-MYC-GFP and Panc28-MYC-GFP-ADAR1 were engineered to express Fcγ receptor 2A fused with the luciferase Luc2 at the C-terminus. A total of 5 × 103 target cells were cultured in a 96-well plate overnight, followed by incubation with 1 × 104 CD8+ T cells in RPMI containing 200 ng/mL OKT3 antibody for 3 days. Next, 20 μg/mL nivolumab, 50 μM ATRA, or a combination were individually added to the mix of target cells and CD8+ T cells. After incubation for 3 days, the 96-well plates were washed with PBS to remove the CD8 T+ cells, and the target cell viability was determined with a CCK-8 kit.

### CCK-8 assay

To quantify cell viability, 3–5 × 103 pancreatic cancer cells were seeded in a 96-well plate. After incubation at 37 °C with 5% CO2 for 1 day, the culture medium was replaced by the indicated concentrations of drugs or CD8+ T cells in 100 μL of the corresponding culture media, and the treated cells were incubated for 3 days. Finally, 20 μL CCK-8 reagent (MedChemExpress) was added into each well, and the optical density at 450 nm was measured using a BioTek Synergy Neo2 plate reader.

### Patients and treatment

A pilot clinical trial was conducted at China Medical University Hospital in Taiwan between March 1, 2021 and February 28, 2022. The study protocol was approved by the Research Ethics Committee, China Medical University & Hospital, Taichung, Taiwan (CMUH110-REC1-013). This trial is registered at ClinicalTrials.gov (NCT05482451). The key inclusion criteria for participants were: age 20 years or older; histologically confirmed pancreatic adenocarcinoma which was locally advanced, recurrent, or metastatic and ineligible or unsuitable for further surgical or radiation interventions; documented disease progression within 6 months after receipt of standard chemotherapies or no available standard chemotherapy; Eastern Cooperative Oncology Group (ECOG) performance status score of 0-2; measurable disease as defined by the Response Evaluation Criteria in Solid Tumors (RECIST) v1.1; and adequate hematologic parameters and hepatic and renal functions. The primary outcome measure was overall response rate. Other endpoints included disease control rate, overall survival, which was calculated from the date of treatment to the death or last follow-up, and safety profiles. All patients had understood and signed informed consent prior to enrollment. All procedures were in accordance with the ethical standards of the responsible committee on human experimentation and with the Declaration of Helsinki of 1975.

Patients received 45 mg/m2 ATRA orally on days 1 to 14 and 3 mg/kg nivolumab intravenously on day 1. The treatment cycle was repeated every 2 weeks. The dose of ATRA was increased to 60 mg/m2 from the second cycle and 75 mg/m2 from the third cycle if patients tolerated the treatment. The treatment continued until disease progression, patient death, intolerable toxicity, withdrawal of informed consent, or at the decision of the in-charge physician.

Physical examination, medical history, ECOG score, and laboratory tests, including blood cell counts, liver function, renal function, biochemistry, and coagulation profiles, were taken at baseline and every 2 weeks. The toxicity profile was graded using the National Cancer Institute Common Terminology Criteria for Adverse Events, version 4.0. Imaging studies using computed tomography were performed at baseline and every 2 months after the initiation of treatment or as clinically indicated. Tumor response was assessed according to RECIST v1.1.

### Colony formation assay

BxPC3, Capan1, and HPAC cells were cultured in 12-well plates containing fresh medium with or without ATRA for 10 to 15 days. The number of colonies with more than 50 cells per colony was counted after staining with crystal violet solution. All the experiments were performed in triplicate wells in 3 independent experiments.

### Immunohistochemical staining of human pancreatic cancer samples

Human tissue microarrays from 135 cases of pancreatic cancer were analyzed by immunohistochemistry staining using ADAR1 antibody (Bethyl Laboratories, Inc.). The study was approved by the Institutional Review Board at The University of Texas MD Anderson Cancer Center. Pancreatic tumor tissues from 9 patients enrolled in the pilot clinical trial were analyzed by immunohistochemical staining using ADAR1 antibody and PD-L1 antibody (Abcam). For PD-L1 staining, the tissue slides were treated with 5% PNGase F containing PBS at 37 °C overnight prior to staining (*40*). Nuclei were counterstained with Mayer’s hematoxylin. The demographics of the 135 pancreatic cancer patients included in the tissue microarrays used for immunohistochemistry are shown in Table S5.

### RNA sequencing and data analysis

Total RNA was isolated using Trizol reagent (Invitrogen). Sequencing libraries were generated using NEBNext Ultra RNA Library Prep Kit for Illumina according to the manufacturer’s instructions. We used HTSeq v0.6.1 to count the number of reads mapped to each gene. Differential expression analysis of 2 groups was performed using the edgeR R package. A corrected P value of 0.05 and an absolute fold change of 1 were considered to indicate significantly differential expression. KEGG pathway analysis (https://www.genome.jp/kegg/) and DisGeNET (https://www.disgenet.org/) analysis were performed on differentially expressed genes.

### Immunoblotting and immunoprecipitation

Equal amounts of protein lysates were separated by sodium dodecyl sulfate-polyacrylamide gel electrophoresis. Separated proteins were electrophoretically transferred to an Immobilon polyvinylidene fluoride membrane rinsed with methanol. The membrane was blocked with PBS containing 5% BSA overnight at 25 °C for 1 to 2 h and then incubated with primary antibodies at 4 °C overnight. After the membranes were washed with PBS-Tween 20, they were incubated with the recommended dilution of secondary antibodies conjugated with horseradish peroxidase (HRP; Jackson ImmunoResearch) at 25 °C for another 1 to 2 h. The protein bands on blots were developed and performed with Immobilon Western Chemiluminescent HRP Substrate (Millipore Corp). Immunoreactive bands were visualized using the ImageQuant LAS 4000 imaging system (GE). ImageJ software was used for densitometry measurements of the Western blots.

For co-immunoprecipitation (co-IP), the lysates were mixed with an equal amount of co-IP buffer (50 mM Tris-HCl (pH 7.4), 150 mM NaCl, 1mM EDTA, 1% NP-40, 1% Na-deoxycholate). The primary antibody or IgG and protein A or G beads (Millipore Corp.) were added to the lysates, and the reactions were incubated at 4 °C at a rotary device for 1 h. The beads were collected by centrifugation and washed gently with co-IP buffer prior to immunoblotting.

### Statistics and reproducibility

The one-tailed independent Student t-test (normal distribution) was used to compare continuous variables between 2 groups. The Pearson correlation test was used for analyzing the correlation between 2 continuous factors. The Kaplan-Meier estimation and the log-rank test were used to compare survival between groups. All data were derived from at least twice independent experiments. Statistical analyses were performed with Microsoft Excel and GraphPad Prism 8. The level of statistical significance was set at P < 0.05 for all tests.

## Acknowledgements

We thank Dr. Zhenbo Han for technical assistance and Dr. Amy Ninetto for editing and proofreading. We thank the MD Anderson Flow Cytometry and Cellular Imaging Core Facility for Mass cytometry (CyTOF) and the MD Anderson Sequencing and Microarray Facility for DNA sequencing. We also thank Novogene Bioinformatics Technology Co., Ltd. For RNA sequencing and analysis.

## Author contributions

C.-F.L. designed and performed the experiments, and analyzed data; Y.W., W.-C.C. and Y.-C.S. performed the experiments and analyzed data; L.-Y.B. and C.-F.C. conducted the pilot clinical trial; J.Y. performed in silico gene expression analysis; H.W. provided tumor tissue arrays. C.-W.C., Y.-Y.C., J.F. and J.L. provides the materials. C.-F.L., L.-Y.B., S.-C.W., and M.-C.H. wrote the manuscript; H.-H.L. designed the experiments and provided intellectual input. R.Y., Y.-N.W., D.Y., P.C., A.M. and H.L. provided scientific inputs; H.Y. and M.-C.H. supervised the study. All authors provided input on the manuscript.

## Data and materials availability

For analyzing RNA sequencing data from published databases, TCGA data for 33 tumor types were obtained from the Broad Institute Firehose website (https://gdac.broadinstitute.org/). CCLE cell line gene expression and drug sensitivity data were obtained from the DepMap data portal (https://depmap.org/portal/download/all/). We used the ISG signature compiled by Liu *et al*. (*13*) to calculate an ISG score for each sample by summation of median absolute deviation–based Z scores of all ISG component genes. Differential *ADAR* mRNA expression in cancer and normal tissue, and correlation of gemcitabine resistance gene signature (GSE6914) and ISG gene signature in tumors were analyzed with GEPIA2 (http://gepia2.cancer-pku.cn/#general). Survival was estimated using KM-Plotter (http://kmplot.com/analysis/). The significant gene sets regulated by ATRA in MDA-MB-231 cells (GSE103426), knockout of *ADAR*-enriched gene sets (GSE122168), gene sets upregulated in gemcitabine-resistant SW1990 cells (GSE79953) and gene sets upregulated in gemcitabine-resistant CFPAC cells (GSE140077) were analyzed using the hallmark gene sets. All the cell lines and the plasmids are available from the authors at request.

## References and Notes

1. Dorothée Duluc et al. Interferon-gamma reverses the immunosuppressive and protumoral properties and prevents the generation of human tumor-associated macrophages. Int J Cancer. 2009 Jul 15;125(2):367–73.

2. J Rodríguez-Villanueva et al. Induction of apoptotic cell death in non-melanoma skin cancer by interferon-alpha. Int J Cancer. 1995 Mar 29;61(1):110–4.

3. C Schindler et al. Interferon-dependent tyrosine phosphorylation of a latent cytoplasmic transcription factor. Science. 1992 Aug 7;257(5071):809–13.

4. Daniel L Barber et al. Restoring function in exhausted CD8 T cells during chronic viral infection. Nature. 2006 Feb 9;439(7077):682–7.

5. Mark J Smyth et al. Type I interferon and cancer immunoediting. Nat Immunol. 2005 Jul;6(7):646–8.

6. Thomas Powles et al. MPDL3280A (anti-PD-L1) treatment leads to clinical activity in metastatic bladder cancer. Nature. 2014 Nov 27;515(7528):558–62.

7. Omid Hamid et al. Safety and tumor responses with lambrolizumab (anti-PD-1) in melanoma. N Engl J Med. 2013 Jul 11;369(2):134–44.

8. Daniel Sanghoon Shin et al. Primary resistance to PD-1 blockade mediated by JAK1/2 mutations. Cancer Discov. 2017 Feb;7(2):188–201.

9. Jianjun Gao et al. Loss of IFN-γ pathway genes in tumor cells as a mechanism of resistance to anti-CTLA-4 therapy. Cell. 2016 Oct 6;167(2):397–404.

10. Herbert A, et al. ADAR and immune silencing in cancer. Trends Cancer. 2019 May;5(5):272–82.

11. Yumeng Wang et al. Systematic characterization of A-to-I RNA editing hotspots in microRNAs across human cancers. Genome Res. 2017 Jul;27(7):1112–1125.

12. Hugh S Gannon et al. Identification of ADAR1 adenosine deaminase dependency in a subset of cancer cells. Nat Commun. 2018 Dec 21;9(1):5450.

13. Huayang Liu et al. Tumor-derived IFN triggers chronic pathway agonism and sensitivity to ADAR loss. Nat Med. 2019 Jan;25(1):95–102.

14. Jeffrey J Ishizuka et al. Loss of ADAR1 in tumours overcomes resistance to immune checkpoint blockade. Nature. 2019 Jan;565(7737):43–48.

15. S Cascinu et al. Pancreatic cancer: ESMO Clinical Practice Guidelines for diagnosis, treatment and follow-up. Ann Oncol. 2010 May;21 Suppl 5:v55–8.

16. J B Patterson et al. Expression and regulation by interferon of a double-stranded-RNA-specific adenosine deaminase from human cells: evidence for two forms of the deaminase. Mol Cell Biol. 1995 Oct;15(10):5376–88.

17. J B Patterson et al. Mechanism of interferon action: double-stranded RNA-specific adenosine deaminase from human cells is inducible by alpha and gamma interferons. Virology. 1995 Jul 10;210(2):508–11.

18. Lieping Chen et al. Anti-PD-1/PD-L1 therapy of human cancer: past, present, and future. J Clin Invest. 2015 Sep;125(9):3384–91.

19. Li-Chuan Chan et al. IL-6/JAK1 pathway drives PD-L1 Y112 phosphorylation to promote cancer immune evasion. J Clin Invest. 2019 Jul 15;129(8):3324–3338.

20. Xiao-Han Tang et al. Retinoids, retinoic acid receptors, and cancer. Annu Rev Pathol. 2011;6:345–64.

21. Riyao Yang et al. Galectin-9 interacts with PD-1 and TIM-3 to regulate T cell death and is a target for cancer immunotherapy. Nat Commun. 2021 Feb 5;12(1):832.

22. Amrutkar M et al. Pancreatic cancer chemoresistance to gemcitabine. Cancers. 2017 Nov 16;9(11).

23. T J Brown et al. Purification and characterization of cytostatic lymphokines produced by activated human T lymphocytes. Synergistic antiproliferative activity of transforming growth factor beta 1, interferon-gamma, and oncostatin M for human melanoma cells. J Immunol. 1987 Nov 1;139(9):2977–83.

24. Gillian I Rice et al. Mutations in ADAR1 cause Aicardi-Goutiè res syndrome associated with a type I interferon signature. Nat Genet. 2012 Nov;44(11):1243–8.

25. Mart M Lamers et al. ADAR1: “Editor-in-chief” of cytoplasmic innate immunity. Front Immunol. 2019 Jul 25;10:1763.

26. Joana M P Desterro et al. SUMO-1 modification alters ADAR1 editing activity. Mol Biol Cell. 2005 Nov;16(11):5115–26.

27. Lemin Li et al. Ubiquitin-dependent turnover of adenosine deaminase acting on RNA 1 (ADAR1) is required for efficient antiviral activity of type I interferon. J Biol Chem. 2016 Nov 25;291(48):24974–24985.

28. Gudas LJ, et al. Retinoids regulate stem cell differentiation. J Cell Physiol. 2011 Feb;226(2):322–30.

29. Larange A, et al. Retinoic acid and retinoic acid receptors as pleiotropic modulators of the immune system. Annu Rev Immunol. 2016 May 20;34:369–94.

30. Tang XH, et al. Retinoids, retinoic acid receptors, and cancer. Annu Rev Pathol. 2011;6:345–64.

31. Antonios Chronopoulos et al. ATRA mechanically reprograms pancreatic stellate cells to suppress matrix remodelling and inhibit cancer cell invasion. Nat Commun. 2016 Sep 7;7:12630.

32. Yasemin Küley-Bagheri, et al. Effects of all-trans retinoic acid (ATRA) in addition to chemotherapy for adults with acute myeloid leukaemia (AML) (non-acute promyelocytic leukaemia (non-APL)). Cochrane Database Syst Rev. 2018 Aug 6;8(8):CD011960.

33. J S Lee et al. Phase I evaluation of all-trans-retinoic acid in adults with solid tumors. J Clin Oncol. 1993 May;11(5):959–66.

34. H C Pitot 4th et al. All-trans retinoic acid: a dose-seeking study in solid tumors. Ann N Y Acad Sci. 1993 Dec 31;691:250–2.

35. Andrea Wang-Gillam et al. Nanoliposomal irinotecan with fluorouracil and folinic acid in metastatic pancreatic cancer after previous gemcitabine-based therapy (NAPOLI-1): a global, randomised, open-label, phase 3 trial. Lancet. 2016 Feb 6;387(10018):545–557.

36. Helmut Oettle et al. Second-line oxaliplatin, folinic acid, and fluorouracil versus folinic acid and fluorouracil alone for gemcitabine-refractory pancreatic cancer: outcomes from the CONKO-003 trial. J Clin Oncol. 2014 Aug 10;32(23):2423–9.

37. Jihoon Kang et al. Nab-paclitaxel plus gemcitabine versus FOLFIRINOX as the first-line chemotherapy for patients with metastatic pancreatic cancer: retrospective analysis. Invest New Drugs. 2018 Aug;36(4):732–741.

38. Richard A Skelton et al. Overcoming the resistance of pancreatic cancer to immune checkpoint inhibitors. J Surg Oncol. 2017 Jul;116(1):55–62.

39. Christine Ferry et al. Cullin 3 mediates SRC-3 ubiquitination and degradation to control the retinoic acid response. Proc Natl Acad Sci U S A. 2011 Dec 20;108(51):20603–8.

40. Heng-Huan Lee et al. Removal of N-linked glycosylation enhances PD-L1 detection and predicts anti-PD-1/PD-L1 therapeutic efficacy. Cancer Cell. 2019 Aug 12;36(2):168–178.e4.

