## Supplementary Figures and Tables for "All-trans retinoic acid-mediated ADAR1 degradation synergizes with PD-1 blockade to suppress pancreatic cancer"

#### Supplementary Figure 1

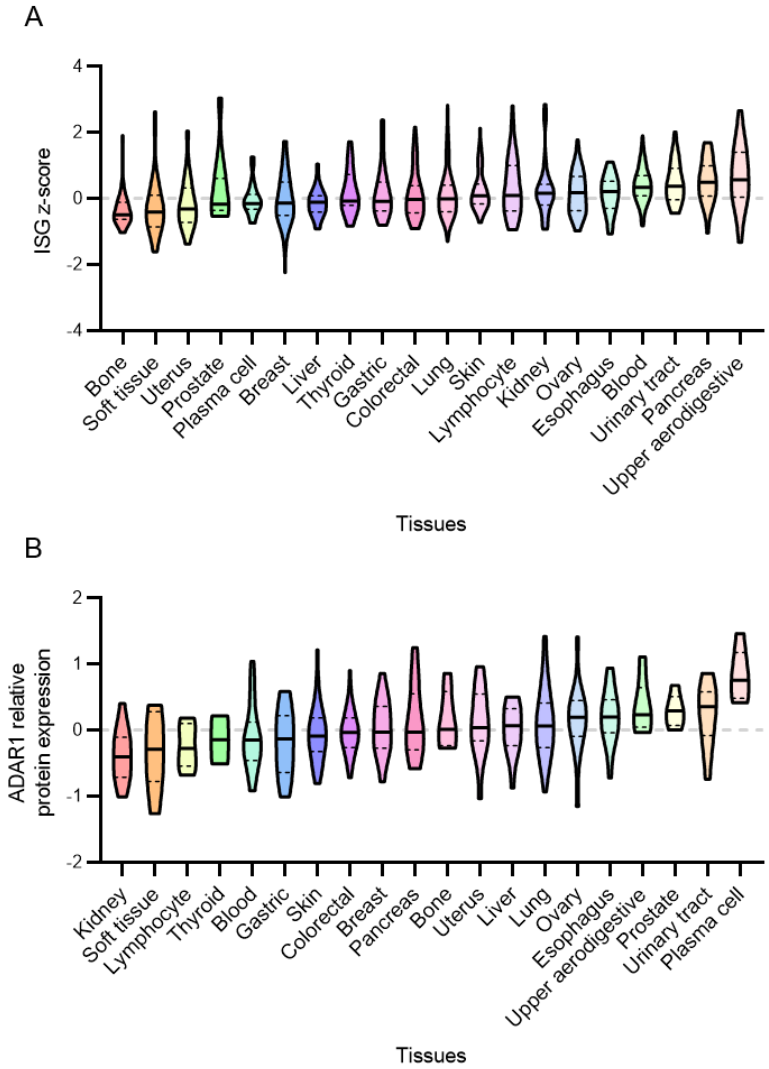


**Figure S1. (A)** Z-scores for interferon-stimulated genes in the indicated tumor types in the Cancer Cell Line Encyclopedia (CCLE). **(B)** ADAR1 relative protein expression in CCLE. Black dotted lines indicate quartiles in each tissue. Black solid lines indicate medians in each tissue.

**Supplementary Figure 2**

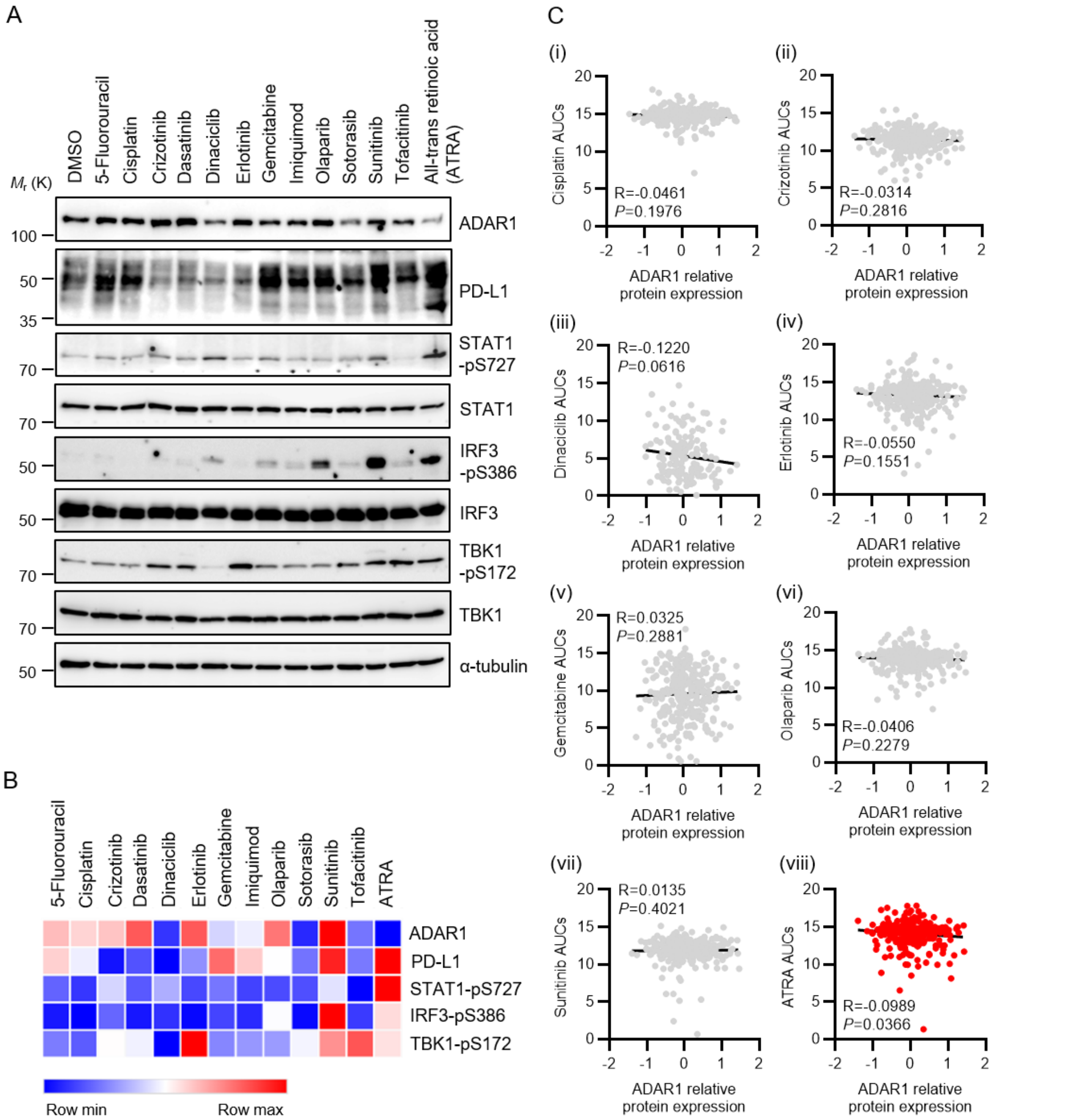


**Figure S2. (A)** Western blots showing the protein levels of ADAR1, PD-L1, STAT1-pS727, IRF3-pS386, and TBK1-pS172 in BxPC3 cells treated with the indicated anticancer drugs for 2 days. Total STAT1, total IRF3, total TBK1 and α-tubulin were loading controls. **(B)** Quantitative proteomic data of ADAR1, PD-L1, STAT1-pS727, IRF3-pS386, and TBK1-pS172 levels in BxPC3 cells visualized by heatmap. **(C)** Pearson correlation analysis of ADAR1 relative protein expression and areas under the receiver operating characteristic curve (AUCs) of (i) cisplatin, (ii) crizotinib, (iii) dasatinib, (iv) erlotinib, (v) gemcitabine, (vi) olaparib, (vii) sunitinib, and (viii) ATRA in the Cancer Cell Line Encyclopedia.

**Supplementary Figure 3**

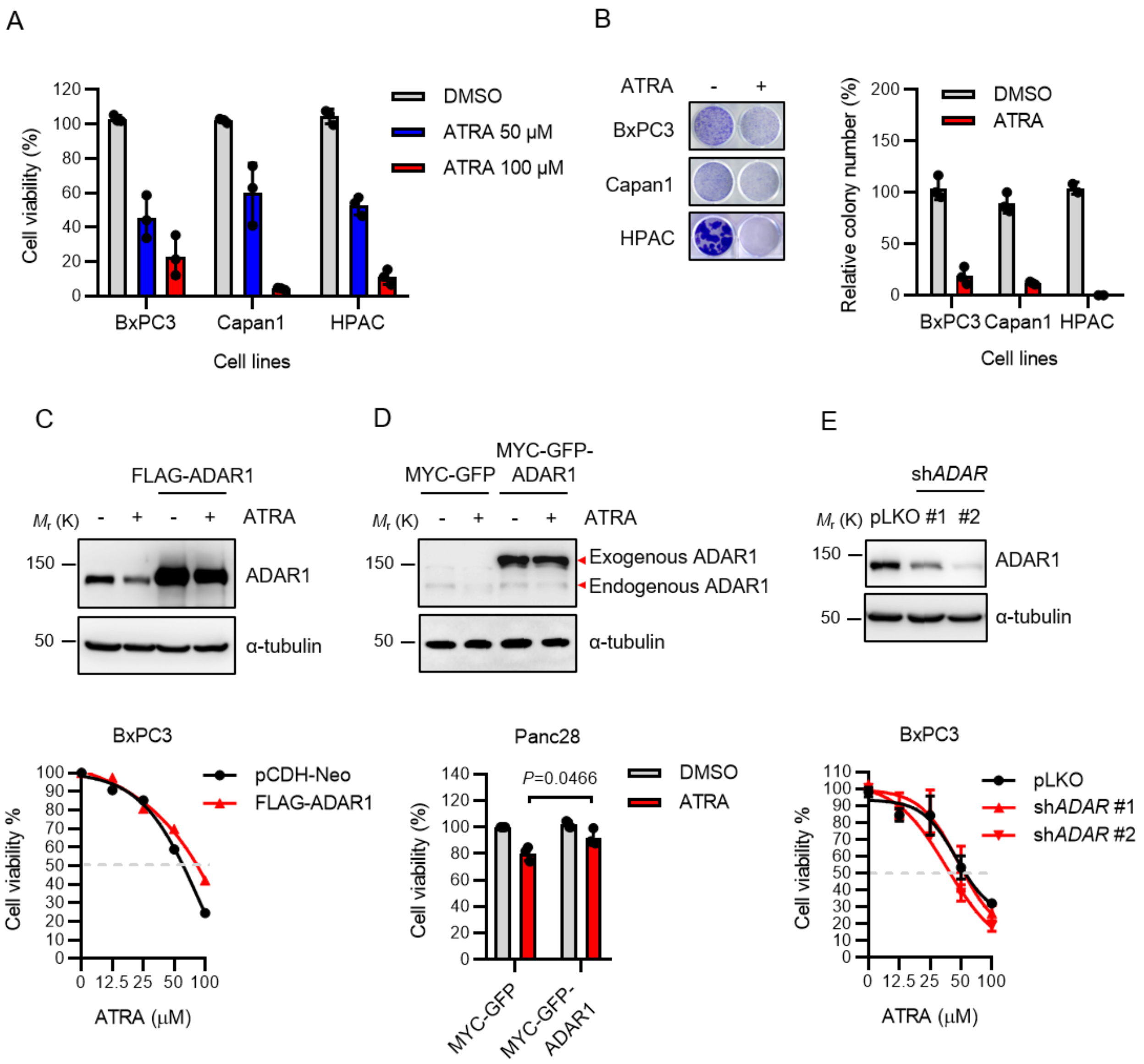


**Figure S3. (A)** CCK-8 assays of BxPC3, Capan1, and HPAC cell viability after treatment with DMSO or ATRA for 3 days. **(B)** Left, representative wells of colony-formation assays of BxPC3, Capan1, and HPAC cells treated with DMSO or 50 µM ATRA for 14 days; right, quantification from three independent experiments performed in duplicate. Data are shown as mean ± SD. *P* values were calculated by Student *t* test. **(C)** Upper, Western blots showing the protein levels of ADAR1 in FLAG-ADAR1 overexpressed BxPC3 cells. Lower, CCK-8 assay of FLAG-ADAR1–overexpressing BxPC3 cell viability after treatment with DMSO or ATRA for 3 days. **(D)** Upper, Western blots showing the protein levels of ADAR1 in MYC-GFP-ADAR1–overexpressing Panc28 cells. Lower, CCK-8 assay of MYC-GFP-ADAR1–overexpressing Panc28 cell viability after treatment with DMSO or ATRA for 3 days. **(e)** Upper, Western blots showing the protein levels of ADAR1 in *ADAR*-knockdown (pLKO) BxPC3 cells. Lower, CCK-8 assay of *ADAR*-knockdown (sh*ADAR*) BxPC3 cell viability after treatment with DMSO or ATRA for 3 days.

**Supplementary Figure 4**

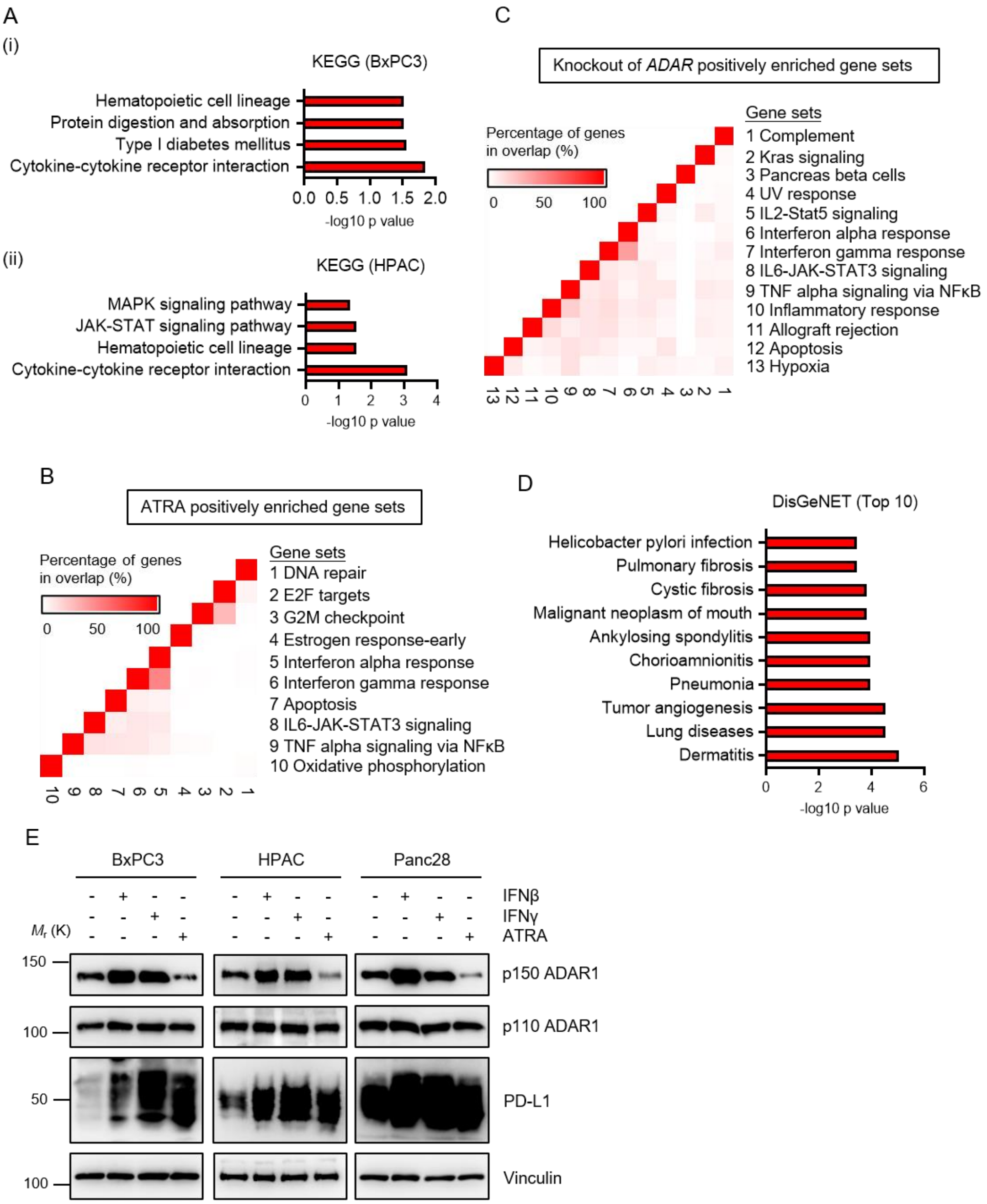


**Figure S4. (A)** Kyoto Encyclopedia of Genes and Genomes (KEGG) pathway enrichment analysis of BxPC3 (i) and HPAC cells (ii) treated with ATRA versus DMSO control. **(B)** Gene Set Enrichment Analysis (GSEA) leading-edge analysis of identified significant gene sets regulated by ATRA in MDA-MB-231 cells (GSE103426). **(C)** GSEA leading-edge analysis of identified significantly enriched gene sets in *ADAR*-knockout H196 cells (GSE122168). **(D)** Top-scored disease associations for ATRA in the DisGeNET database. **(E)** Western blots showing the protein levels of ADAR1 p110 isoform, ADAR1 p150 isoform, and PD-L1 in BxPC3, HPAC, and Panc28 cells treated with 20 ng/mL interferon (IFN)-β, 20 ng/mL IFN-γ, or 50 μM ATRA for 24 h.

#### Supplementary Figure 5

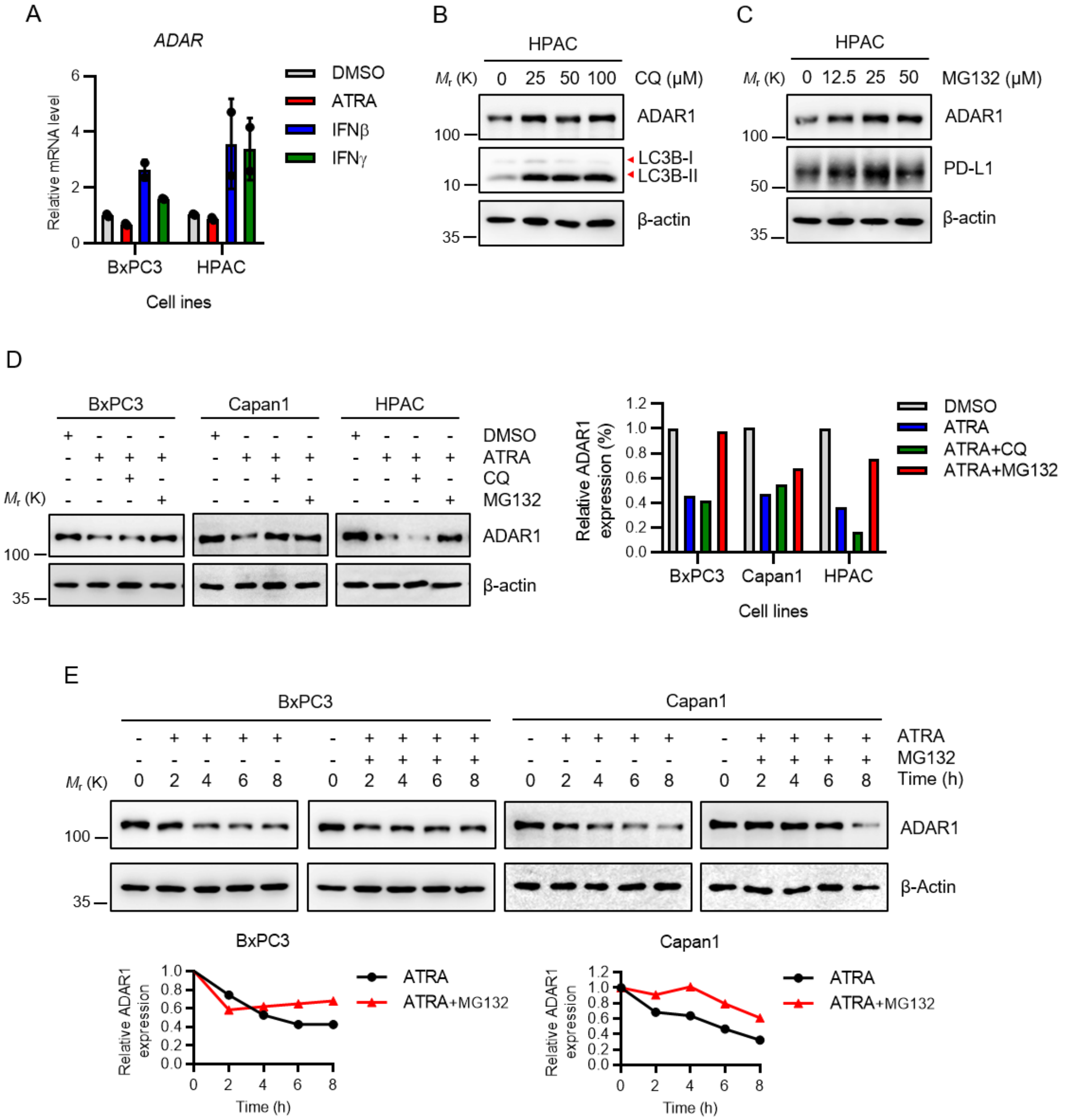


**Figure S5. (A)** Expression of *ADAR* in BxPC3 and HPAC cells treated with 20 ng/mL IFNβ, 20 ng/mL IFNγ, or 50 μM ATRA for 24 h. Data are shown as mean ± SD. *P* values were calculated by Student *t* test. **(B)** Western blots showing the protein levels of ADAR1 in HPAC cells treated with the indicated concentrations of chloroquine (CQ) for 24 h. **(C)** Western blots showing the protein levels of ADAR1 in HPAC cells treated with the indicated concentrations of MG132 for 24 h. **(D)** Left, Western blots for showing the protein levels of ADAR1 in BxPC3, Capan1, and HPAC cells treated with ATRA, CQ, or MG132 for 24 h. Right, quantitative ADAR1 expression. **(E)** Upper, Western blots showing the protein levels of ADAR1 in BxPC3 and Capan1 cells treated with ATRA and MG132 alone or combination for the indicated time. Lower, quantitative ADAR1 expression.

**Supplementary Figure 6**

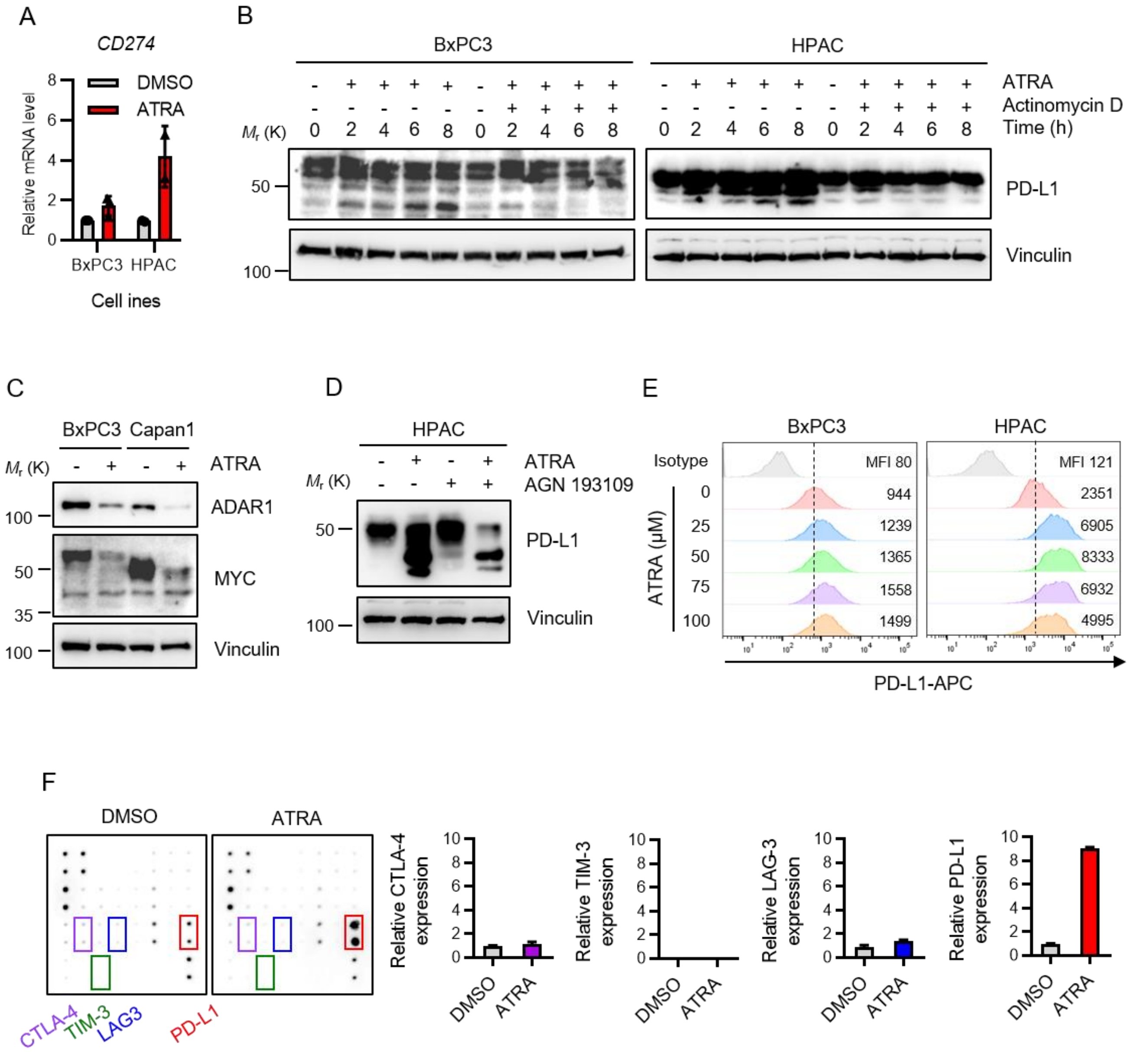


**Figure S6. (A)** Expression of *CD274* gene (PD-L1) in BxPC3 and HPAC cells treated with or without 50 μM ATRA for 24 h. Data represent mean ± SD. *P* values were calculated by Student *t* test. **(B)** Western blots showing the protein levels of PD-L1 in BxPC3 and HPAC cells treated with ATRA and actinomycin D alone or in combination for the indicated time. **(C)** Western blots showing exogenous MYC-tagged PD-L1 protein expression in BxPC3 and Capan1 cells treated with ATRA for 24 h. **(D)** Western blots for showing the protein levels of PD-L1 in HPAC cells treated with 50 μM ATRA or 10 μM AGN 193109 for 24 h. **(E)** Flow cytometry analysis showing surface PD-L1 expression in BxPC3 and HPAC cells treated with the indicated concentrations of ATRA for 24 h. **(F)** Left, immune checkpoint arrays developed using lysates from HPAC cells treated with or without ATRA. Right, quantitative LAG-3, TIM-3, CTLA-4, and PD-L1 expression.

**Supplementary Figure 7**

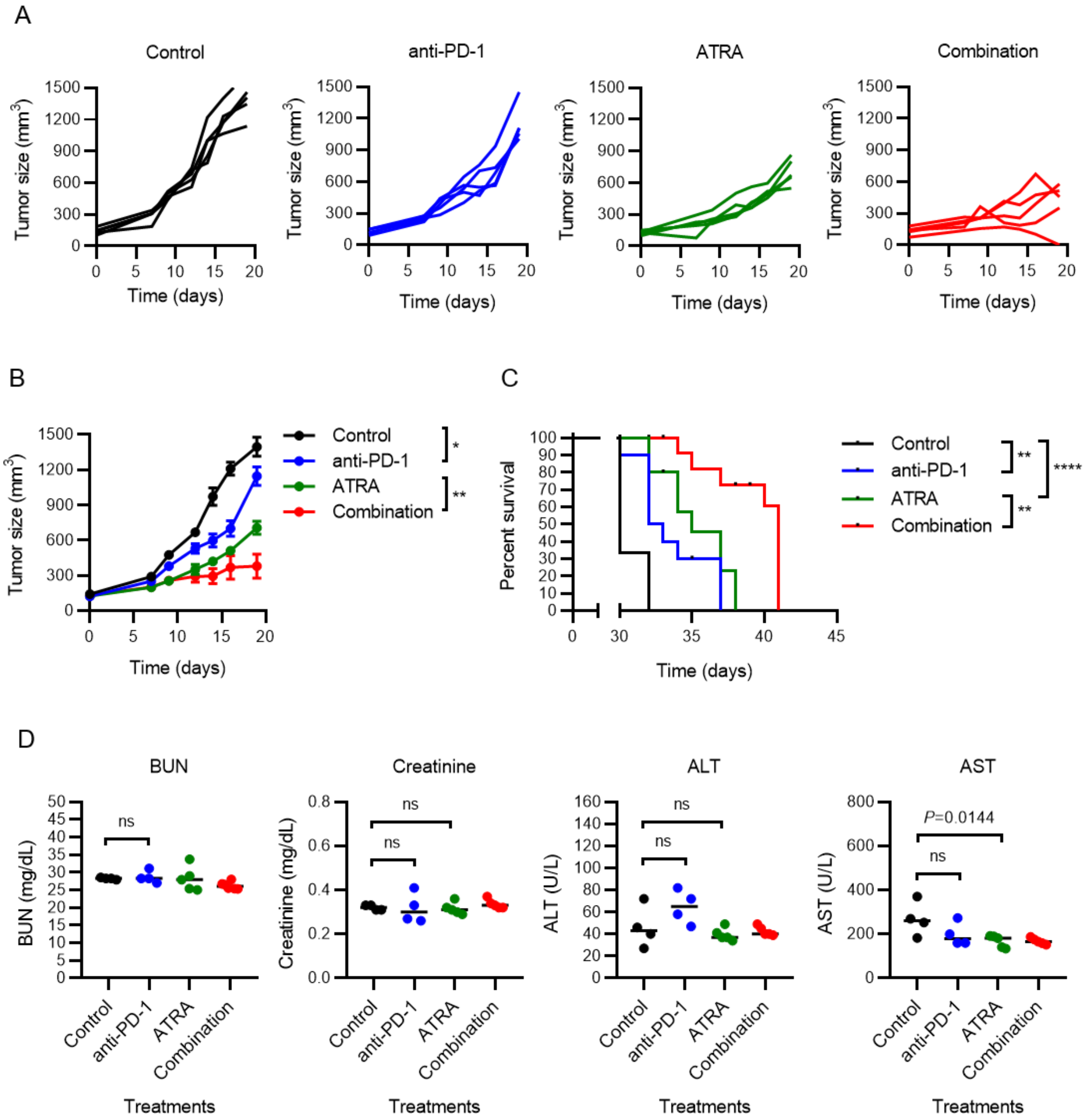


**Figure S7. (A)** Tumor growth curves of individual BALB/cJ mice orthotopically injected with 4T1 tumor cells and subjected to the indicated treatments. **(B)** Mean tumor size in the mice orthotopically injected with 4T1 tumor cells and subjected to the indicated treatments. Dots represent mean (5 mice/group); whiskers represent SD. **(C)** Comparison of Kaplan-Meier survival curves (log-rank tests) of 4T1 tumor–bearing mice. **(D)** Blood urea nitrogen (BUN), creatinine, alanine aminotransferase (ALT), and aspartate aminotransferase (AST) levels in blood plasma of treated C57BL/6 mice bearing Panc02 cells. Each dot represents individual values. The black bars represent the mean values. *P* values were calculated by Student *t* test. **P* < 0.05; ***P* < 0.01; ****P* < 0.001; *****P* < 0.0001; ns, not significant.

**Supplementary Figure 8**

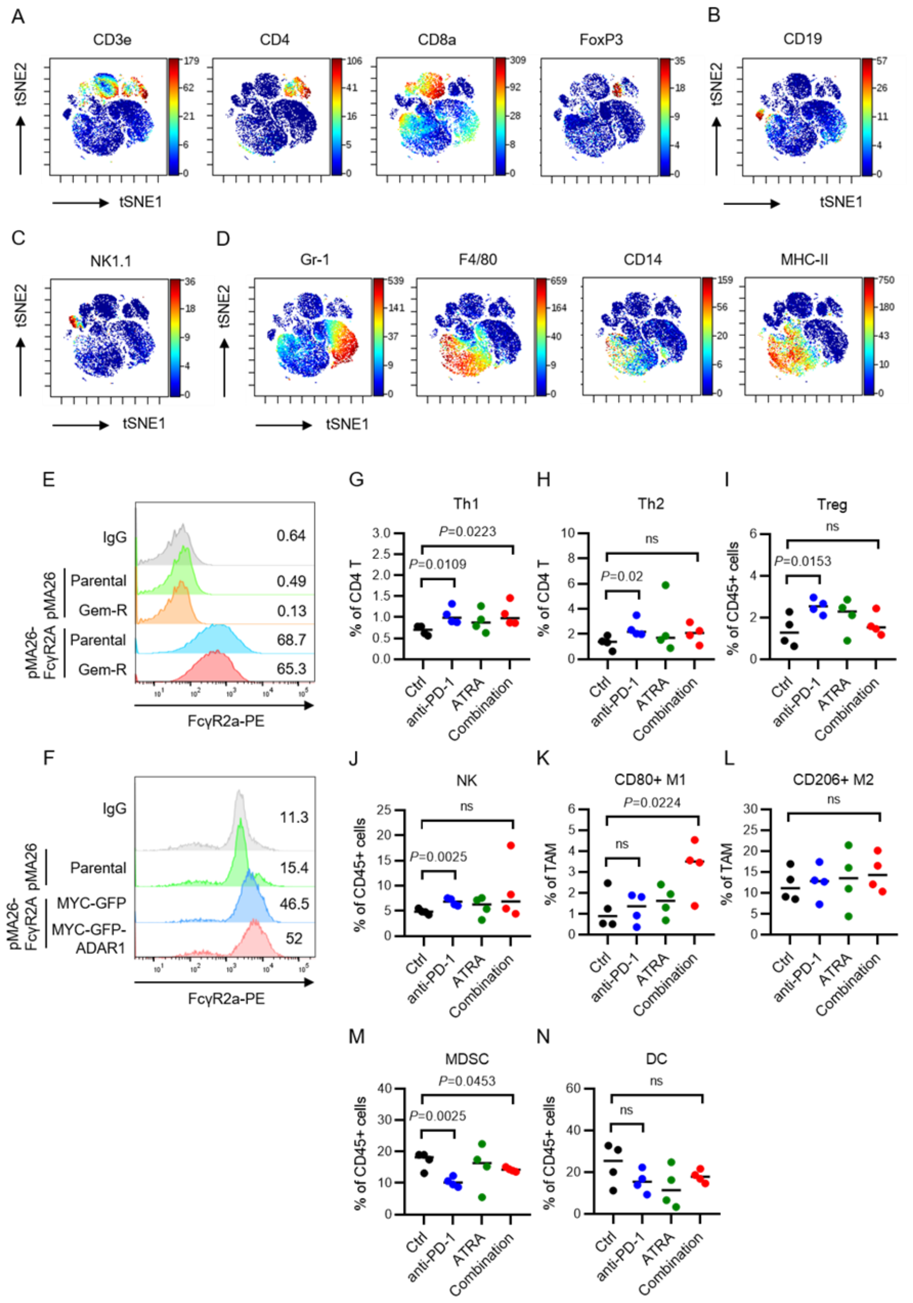


**Figure S8.** **(A)** T-distributed stochastic neighbor embedding (t-SNE) plots of tumor-infiltrating lymphocytes (TILs) overlaid with the expression of selected markers for T cells, **(B)** B cells, **(C)** natural killer (NK) cells, and **(D)** myeloid cells. **(E)** Flow cytometry analysis showing surface Fcγ receptor 2A expression in Panc28-RL2 and Panc28-Gem-R-RL2 cells. **(F)** Flow cytometry analysis showing the surface Fcγ receptor 2A expression in Panc28-RL2-MYC-GFP and Panc28-RL2-MYC-GFP-ADAR1 cells. Time-of-flight mass cytometry analysis of T helper 1 (Th1) **(G)**, Th2 **(H)**, regulatory T (Treg) **(I)**, NK **(J)**, CD80+ M1 macrophage **(K)**, CD206+ M2 macrophage **(L)**, myeloid-derived suppressor cell (MDSC) **(M)**, and dendritic cell (DC) **(N)** immune cell populations in TILs. Each dot represents individual values. The black bars represent the mean values. *P* values were calculated by Student *t* test. Gem-R, gemcitabine-resistant.

**Supplementary Figure 9**

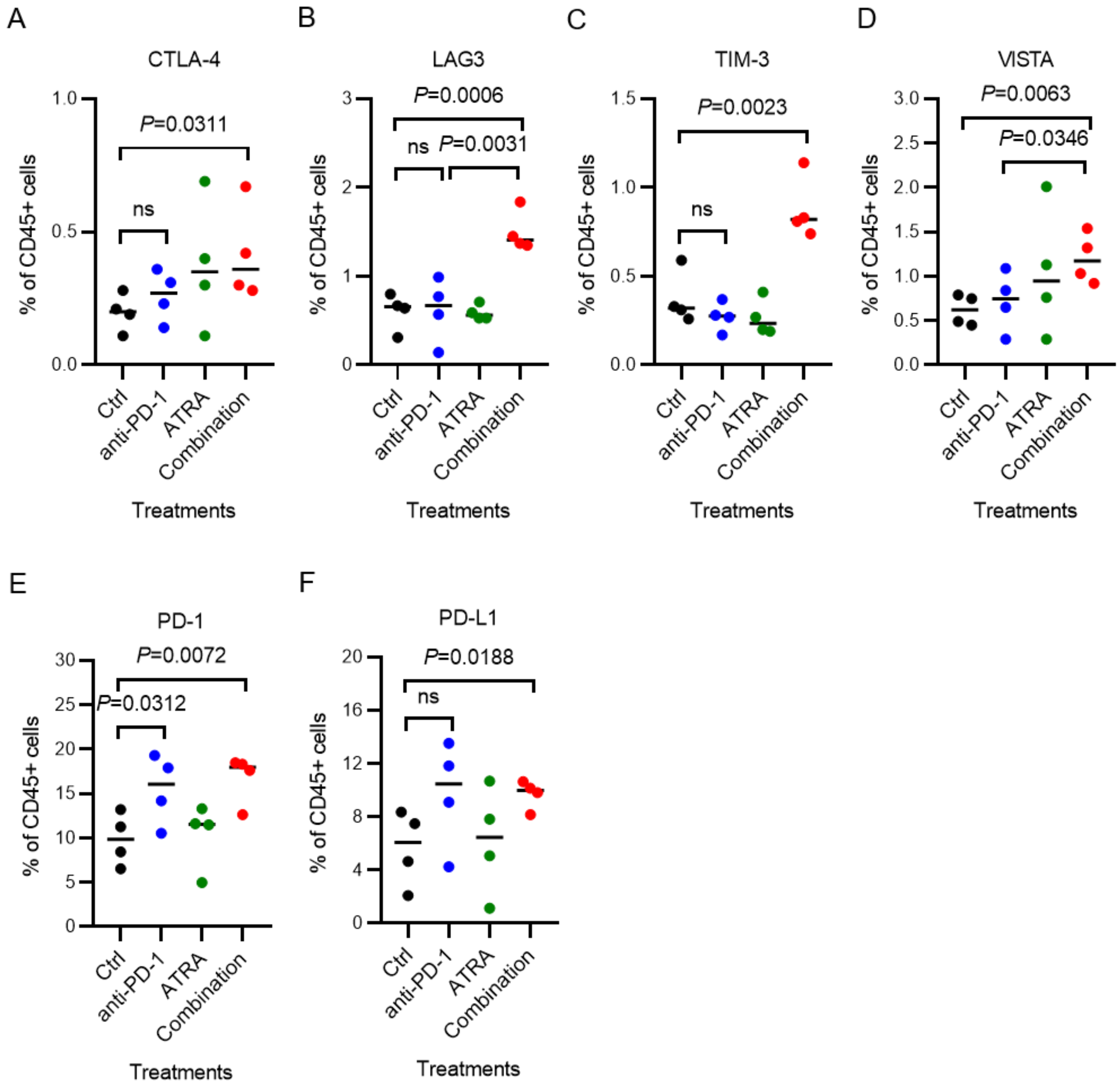


**Figure S9.** Time-of-flight mass cytometry analysis of co-expression of co-inhibitory receptors CTLA-4 **(A)**, LAG3 **(B)**, TIM-3 **(C)**, VISTA **(D)**, PD-1 **(E),** and PD-L1 **(F)** in tumor-infiltrating lymphocytes under the indicated treatments. *P* values were calculated using Student *t* test. Dots indicate individual values; black bars indicate mean. ns, not significant.

**Supplementary Figure 10**

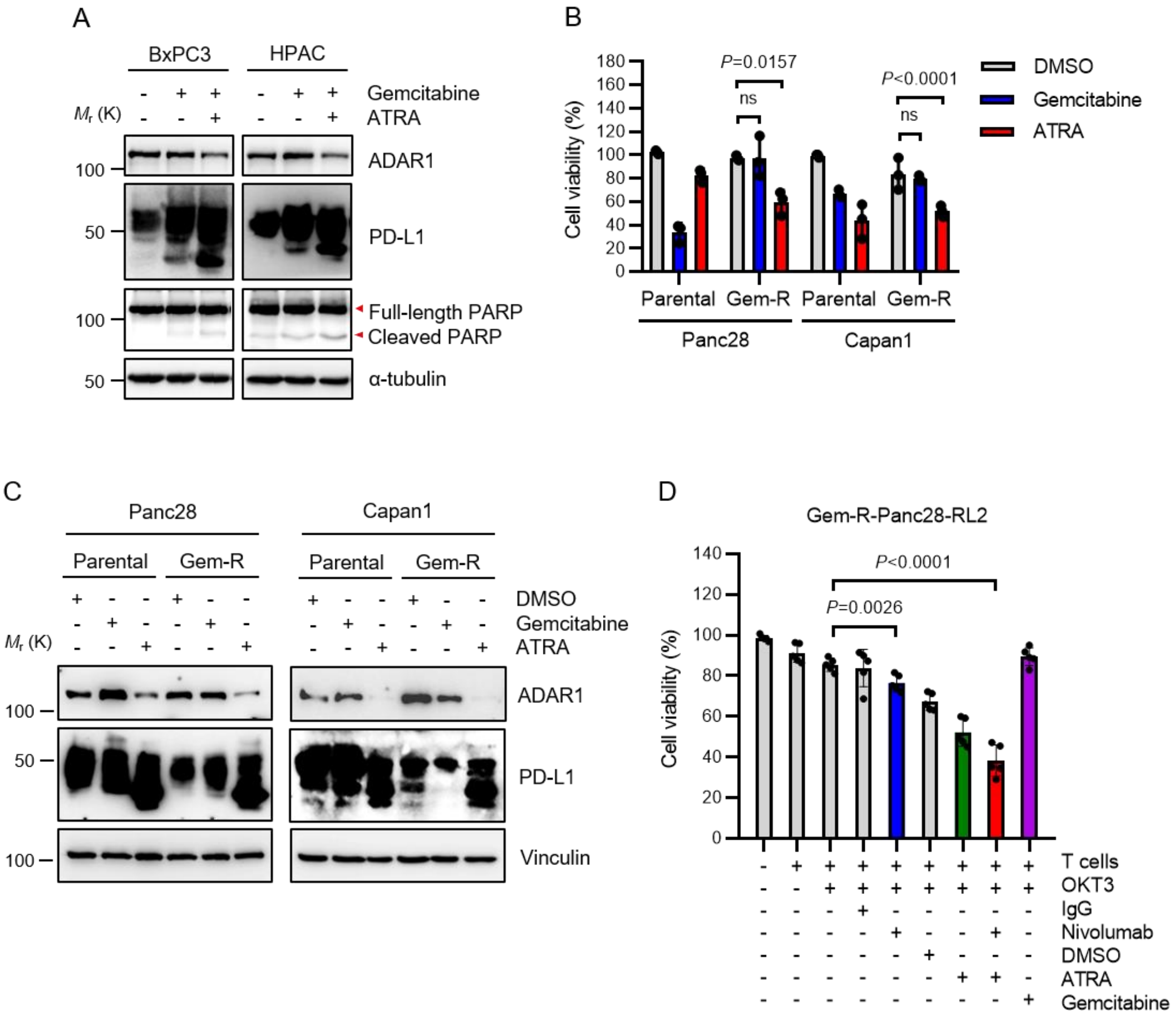


**Figure S10. (A)** Western blots showing the protein levels of ADAR1, PD-L1, and PARP1 in BxPC3 and HPAC cells treated with 1 μM gemcitabine with or without 50 μM ATRA for 24 h. **(B)** CCK-8 assay of parental and gemcitabine-resistant (Gem-R) Capan1 cell viability after treatment with DMSO or ATRA for 3 days. **(C)** Western blots showing the protein levels of ADAR1 and PD-L1 in parental and Gem-R cells treated with 1 μM gemcitabine or 50 μM ATRA for 24 h. **(D)** T-cell cytotoxicity (CCK-8) assays of potency of nivolumab, ATRA, nivolumab + ATRA, and gemcitabine. Target cells were Gem-R-Panc28-RL2 cells. Data represent mean ± SD. *P* values were calculated by Student *t* test. ns, not significant.

#### Supplementary Figure 11

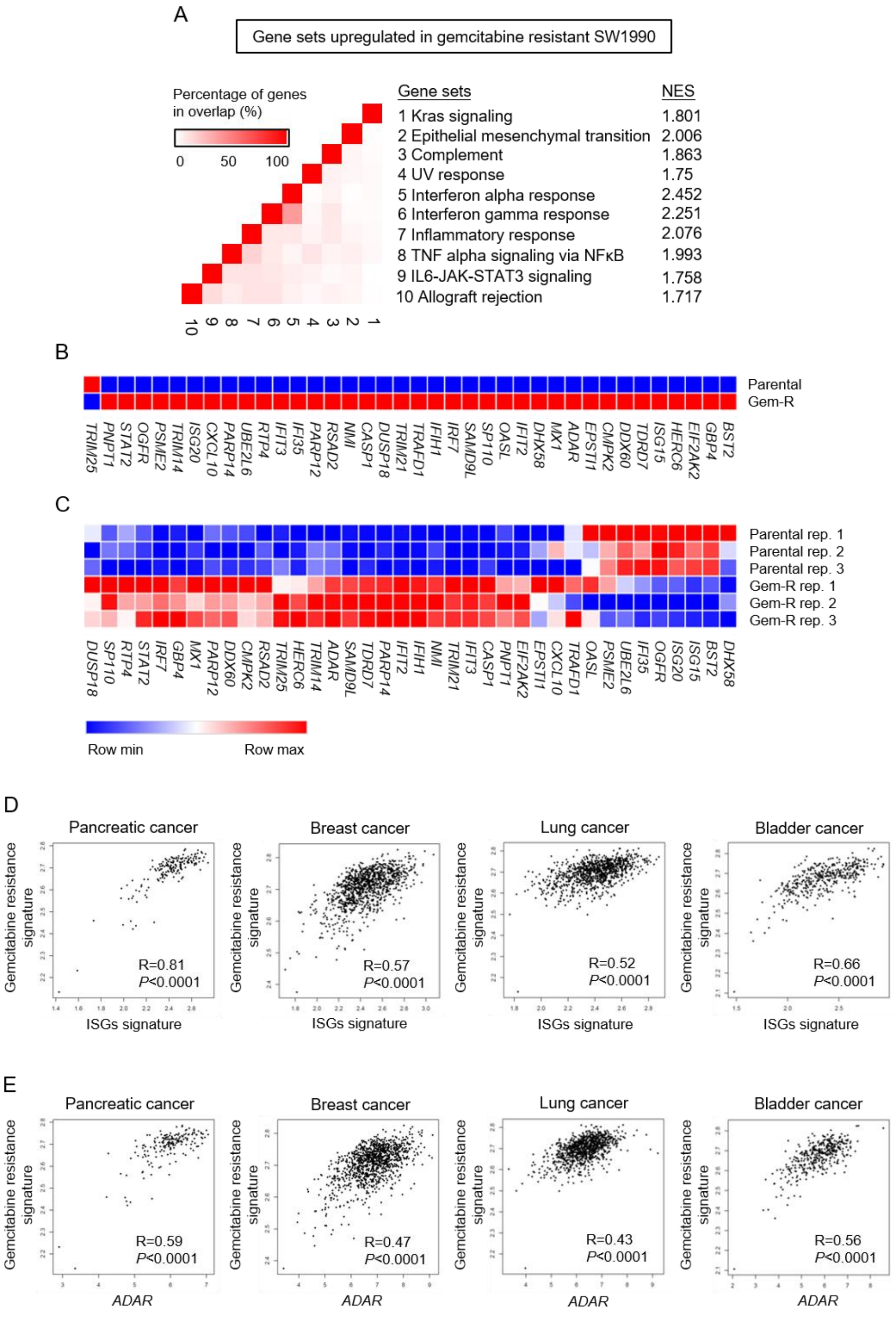


**Figure. S11. (A)** GSEA leading-edge analysis of identified significant gene sets regulated by ATRA in SW1990 pancreatic cancer cells (GSE79953). **(B)** Heatmaps of IFN-stimulated genes (ISGs) in SW1990 cells treated with or without ATRA (GSE79953). **(C)** The heatmaps of ISGs in parental and gemcitabine-resistant (Gem-R) CFPAC pancreatic cancer cells (GSE140077). **(D)** Pearson correlation analysis of gemcitabine resistance gene signature (GSE6914) and ISGs signature of TCGA pancreatic cancer, breast cancer, lung cancer and bladder cancer datasets. **(E)** Pearson correlation analysis of gemcitabine resistance gene signature (GSE6914) and *ADAR* expression of TCGA pancreatic cancer, breast cancer, lung cancer and bladder cancer datasets.

### Supplementary Tables

***Table S1. Characteristics of 9 PDAC patients for the pilot clinical trial***

| | | | | | | | | | |
|---|---|---|---|---|---|---|---|---|---|
| Subject number | 1 | 2 | 3 | 4 | 5 | 6 | 7 | 8 | 9 |
| Age (years) | 63 | 79 | 78 | 55 | 54 | 68 | 57 | 69 | 61 |
| Gender | M | M | M | F | F | F | F | F | M |
| Number of lines for prior chemotherapy | 3 | 4 | 2 | 2 | 2 | 3 | 2 | 3 | 4 |
| Highest dose of ATRA ($mg/m^2$/day) | 45 | 45 | 60 | 45 | 45 | 75 | 75 | 75 | 60 |
| Completed cycles | 4 | 4 | 8 | 1 | 4 | 11 | 10 | 4 | 3 |
| Best of response | PD | PD | SD | PD | PD | SD | SD | PD | PD |
| Overall survival (week) | 12.571 | 10 | 23.429 | 3.143 | 6.857 | 21.857 | 23.143 | 15.571 | 9 |

Abbreviation: F, female; M, male; PD, progressive disease; SD, stable disease

***Table S2. CyTOF antibody list***

| Target | Label | Intracellular staining | Clone | Source | Catalog number |
|---|---|---|---|---|---|
| CD45(Ms) | 89Y | FALSE | 30-F11 | DVS-Fluidigm | 3089005B |
| CD4(Ms) | 115In | FALSE | RM4-5 | BioLegend | 100506 |
| Ly-6G/Ly-6C, Gr-1 | 141Pr | FALSE | RB6-8C5 | BioLegend | 108402 |
| CD11c | 142Nd | FALSE | N418 | DVS-Fluidigm | 3142003B |
| GATA3 | 145Nd | TRUE | TWAJ | eBioscience | 14-9966-82 |
| CD8a | 146Nd | FALSE | 53-6.7 | BioLegend | 100702 |
| CD223, LAG-3 | 147Sm | FALSE | C9B7W | BioLegend | 125202 |
| CD11b | 148Nd | FALSE | M1/70 | DVS-Fluidigm | 3148003B |
| CD19 | 149Sm | FALSE | 4D5 | BioLegend | 115502 |
| CD25 | 150Nd | FALSE | 3C7 | BioLegend | 101902 |
| CD3, CD3e | 152Sm | FALSE | 145-2C11 | BioLegend | 100302 |
| Arginase 1 | 153Eu | TRUE | Polyclonal | R&D | AF5868 |
| CD274, PD-L1 | 154Sm | FALSE | 10F.9G2 | BioLegend | 124303 |
| CD14 | 156Gd | FALSE | Sa14-2 | DVS-Fluidigm | 3156009B |
| Foxp3 | 158Gd | TRUE | FJK-16s | DVS-Fluidigm | 3158003A |
| CD279, PD-1 | 159Tb | FALSE | 29F.1A12 | BioLegend | 135202 |
| VISTA, B7-H5 | 160Gd | FALSE | BLR035F | BETHYL | A700-035CF |
| iNOS | 161Dy | TRUE | CXNFT | DVS-Fluidigm | 3161011B |
| TIM3, CD366 | 162Dy | FALSE | RMT3-23 | DVS-Fluidigm | 3162029B |
| CD152, CTLA-4 | 163Dy | FALSE | 9H10 | BioLegend | 106202 |
| IFNg | 165Ho | TRUE | XMG1.2 | DVS-Fluidigm | 3165003B |
| IL-4 | 166Er | TRUE | 11B11 | DVS-Fluidigm | 3166003B |
| CD206, MMR | 169Tm | TRUE | C068C2 | DVS-Fluidigm | 3169021B |
| NK1.1 | 170Er | FALSE | PK136 | BioLegend | 108702 |
| CD80 | 171Yb | FALSE | 16-10A10 | DVS-Fluidigm | 3171008B |
| CD86 | 172Yb | FALSE | GL1 | DVS-Fluidigm | 3172016B |
| F4/80 | 173Yb | FALSE | BM8 | BioLegend | 123102 |
| IL-17A | 174Yb | TRUE | TC11-18H10.1 | DVS-Fluidigm | 3174002B |
| I-A/I-E, MHC-II | 209Bi | FALSE | M5/114.15.2 | DVS-Fluidigm | 3209006B |

***Table S3. The markers for identifying major TIL subsets***

| Population | Marker |
|---|---|
| CD4 T | CD45+CD11b-CD3+CD8-CD4+ |
| Th1 | CD45+CD11b-CD3+CD8-CD4+IFNr+ |
| Th2 | CD45+CD11b-CD3+CD8-CD4+GATA-3+IL4+ |
| CD8 T | CD45+CD11b-CD3+CD8+CD4- |
| Treg | CD45+CD11b-CD3+CD8-CD4+Foxp3+CD25+ |
| MDSC | CD45+CD11b+CD3-Gr1+ |
| Monocyte | CD45+CD11b+CD3-Gr1-F4/80-CD14+ |
| TAM | CD45+CD11b+CD3-Gr1-F4/80+ |
| M1 | CD45+CD11b+CD3-Gr1-F4/80+CD80+ |
| M2 | CD45+CD11b+CD3-Gr1-F4/80+CD206+ |
| B | CD45+CD11b+CD3-CD19+ |
| NK | CD45+CD11b+CD3-NK1.1+ |
| DC | CD45+CD11b+CD3-MHC-II+CD11c+ |

Abbreviation: Th1, T helper 1; Th2, T helper 2; Treg, regulatory T; MDSC, myeloid-derived suppressor cell; TAM, Tumor-associated macrophage; NK, nature killer; DC, dendritic cell

***Table S4. Information of primer sequences for RT-qPCR***

| Gene | Forward | Reverse |
|---|---|---|
| *ADAR* | CATGGCTTTGCTGCTGAAT | CTGCTTGCCTTGCTTCTTG |
| *CD274* | GGACAAGCAGTGACCATCAAG | CCCAGAATTACCAAGTGAGTCCT |
| *GAPDH* | GGAGCGAGATCCCTCCAAAAT | GGCTGTTGTCATACTTCTCATGG |

*Table S5. Characteristics of 135 PDAC patients for IHC of ADAR1*

| Subject number | Gender | Differentiation | AJCC Stage | Survival (days) |
|---|---|---|---|---|
| 1 | F | Poor | IIB | 13.38 |
| 2 | M | Moderate | IIB | 9.90 |
| 3 | F | Moderate | IIA | 9.99 |
| 4 | M | Moderate | IIA | 7.53 |
| 5 | M | Poor | IV | 5.82 |
| 6 | F | Moderate | IIB | 5.23 |
| 7 | F | Moderate | IV | 15.55 |
| 8 | M | Well | IIB | 42.64 |
| 9 | F | Moderate | IIA | 9.60 |
| 10 | M | Poor | IIB | 21.21 |
| 11 | M | Mod | IIA | 12.69 |
| 12 | M | Moderate | IIB | 29.36 |
| 13 | M | Poor | IV | 8.12 |
| 14 | F | Poor | IIA | 17.95 |
| 15 | M | Moderate | IIB | 10.45 |
| 16 | F | Poor | IIA | 128.84 |
| 17 | F | Moderate | IIB | 78.38 |
| 18 | M | Moderate | IIB | 10.39 |
| 19 | M | Poor | IIB | 44.25 |
| 20 | M | Poor | IIA | 7.69 |
| 21 | M | Moderate | IIB | 10.39 |
| 22 | M | Moderate | IIB | 36.23 |
| 23 | F | Moderate-Poor | IIB | 17.69 |
| 24 | M | Moderate | IIB | 18.94 |
| 25 | M | Moderate | IIB | 40.60 |
| 26 | F | Moderate | IIB | 26.53 |
| 27 | F | Mod-Poor | IIB | 14.04 |
| 28 | M | Poor | IIA | 105.67 |
| 29 | M | Mod | IIB | 24.99 |
| 30 | F | Mod-poor | IIB | 12.72 |
| 31 | M | Moderate-Poor | IIB | 43.20 |

***Table S5. Characteristics of 135 PDAC patients for IHC of ADAR1 (continued)***

| Subject number | Gender | Differentiation | AJCC Stage | Survival (days) |
|---|---|---|---|---|
| 32 | M | Mod | IIB | 37.12 |
| 33 | M | Moderate | IIB | 74.83 |
| 34 | F | Mod | IIB | 11.31 |
| 35 | F | Mod | IIB | 21.83 |
| 36 | M | Mod | IIB | 22.52 |
| 37 | M | Mod-poor | IIB | 5.42 |
| 38 | M | Mod | IIB | 51.68 |
| 39 | F | Mod | IIB | 17.75 |
| 40 | F | Mod-poor | IIB | 9.04 |
| 41 | M | Mod-Poor | IIA | 11.70 |
| 42 | M | Mod | IIB | 16.87 |
| 43 | F | Moderate-Poor | IIA | 96.26 |
| 44 | F | Moderate | IIB | 38.27 |
| 45 | F | Mod | IIB | 34.49 |
| 46 | F | Mod-poor | IIB | 7.63 |
| 47 | M | Mod-poor | IIB | 20.25 |
| 48 | F | Mod | IIB | 12.26 |
| 49 | M | Mod-poor | III | 33.53 |
| 50 | F | Mod | IIB | 14.63 |
| 51 | M | Poor | IIB | 25.22 |
| 52 | M | Poor | IIB | 76.44 |
| 53 | F | Poor | IIA | 18.21 |
| 54 | F | Moderate | IIB | 10.16 |
| 55 | F | Poor | IIB | 69.50 |
| 56 | M | Mod | IIB | 31.63 |
| 57 | M | Mod | IIB | 55.99 |
| 58 | F | Mod | IIB | 18.51 |
| 59 | M | Mod | IIB | 12.10 |
| 60 | F | Mod | IIB | 32.71 |

***Table S5. Characteristics of 135 PDAC patients for IHC of ADAR1 (continued)***

| Subject number | Gender | Differentiation | AJCC Stage | Survival (days) |
|---|---|---|---|---|
| 61 | M | Moderate | IIB | 78.05 |
| 62 | F | Poor | IIB | 5.29 |
| 63 | F | Mod | IIB | 70.95 |
| 64 | F | Mod | IIB | 20.19 |
| 65 | M | Mod | IIB | 4.18 |
| 66 | M | Mod | IIB | 21.21 |
| 67 | F | Mod | IIB | 25.32 |
| 68 | M | Mod | IIA | 44.12 |
| 69 | F | Mod | IIB | 4.67 |
| 70 | F | Mod | IIB | 12.49 |
| 71 | M | Mod | IIB | 15.88 |
| 72 | F | Mod | IIB | 33.14 |
| 73 | F | Mod | IIB | 10.65 |
| 74 | M | Mod-poor | IIB | 8.22 |
| 75 | M | Mod | IIA | 16.80 |
| 76 | M | Mod | IIB | 61.38 |
| 77 | M | Mod | IIB | 25.41 |
| 78 | F | Mod | IIB | 26.89 |
| 79 | M | Mod | IIB | 51.02 |
| 80 | F | Mod | IIB | 60.59 |
| 81 | M | Mod | IIB | 0.43 |
| 82 | M | Mod | IIB | 56.75 |
| 83 | F | Mod | IIB | 7.69 |
| 84 | M | Mod | IIB | 35.05 |
| 85 | M | Poor | IIB | 11.11 |
| 86 | M | Mod | IIA | 41.56 |
| 87 | F | Mod | IIB | 32.22 |
| 88 | M | Poor | IIB | 41.19 |
| 89 | F | Mod | IIA | 29.42 |

***Table S5. Characteristics of 135 PDAC patients for IHC of ADAR1 (continued)***

| Subject number | Gender | Differentiation | AJCC Stage | Survival (days) |
|---|---|---|---|---|
| 90 | M | Mod | IIB | 41.16 |
| 91 | F | Mod | IIB | 35.90 |
| 92 | F | Mod | IIB | 17.19 |
| 93 | M | Poor | IIB | 18.38 |
| 94 | F | Mod | IIB | 9.96 |
| 95 | M | Poor | IIB | 21.76 |
| 96 | F | Poor | IIB | 7.79 |
| 97 | M | Poor | IIA | 29.65 |
| 98 | F | Poor | IIB | 27.58 |
| 99 | M | Poor | IIB | 32.81 |
| 100 | F | Mod | IIB | 25.68 |
| 101 | F | Mod | IB | 26.27 |
| 102 | F | Mod | IIB | 13.41 |
| 103 | M | Mod | IIA | 21.76 |
| 104 | M | Moderate | IIA | 14.43 |
| 105 | F | Poor | IIB | 25.02 |
| 106 | M | Moderate | IIA | 17.69 |
| 107 | F | Moderate | IIB | 12.69 |
| 108 | M | Moderate | IIB | 13.28 |
| 109 | M | Poor | IIB | 236.52 |
| 110 | M | Well | IIA | 23.38 |
| 111 | M | Moderate | IIB | 13.45 |
| 112 | F | Moderate | IIA | 4.18 |
| 113 | M | Moderate | IIB | 22.39 |
| 114 | F | Moderate | IIA | 82.88 |
| 115 | F | Poor | IIB | 4.34 |
| 116 | F | Moderate | IIB | 120.39 |
| 117 | M | Moderate | IIB | 72.20 |
| 118 | M | Well | IIB | 154.03 |

***Table S5. Characteristics of 135 PDAC patients for IHC of ADAR1 (continued)***

| Subject number | Gender | Differentiation | AJCC Stage | Survival (days) |
|---|---|---|---|---|
| 119 | F | Moderate | IIB | 11.67 |
| 120 | M | Poor | IIB | 6.18 |
| 121 | M | Poor w SRC | IIB | 67.76 |
| 122 | M | Moderate | IIA | 236.25 |
| 123 | F | Well | IIA | 137.26 |
| 124 | M | Well | IIB | 33.44 |
| 125 | M | Moderate | IIB | 20.94 |
| 126 | M | Moderate | IIB | 16.04 |
| 127 | M | Moderate | IIA | 9.50 |
| 128 | M | Moderate | IIA | 18.74 |
| 129 | M | Moderate | IIB | 19.46 |
| 130 | F | Moderate | IIB | 10.75 |
| 131 | M | Moderate | IIA | 207.12 |
| 132 | M | Poor | IIB | 14.60 |
| 133 | F | Moderate | IIA | 65.16 |
| 134 | M | Moderate | IIA | 63.22 |
| 135 | M | Moderate | IIA | 38.86 |

Abbreviation: AJCC, the American Joint Committee on Cancer
